# Structural insights into the heterotrimeric alternatively spliced P2X7 receptors

**DOI:** 10.1101/2023.05.30.542804

**Authors:** Sophie K. F. De Salis, Jake Zheng Chen, Kristen K. Skarratt, Stephen J. Fuller, Thomas Balle

## Abstract

AlphaFold2-Multimer was used to generate structures of the heterotrimeric P2X7 receptors composed of wild-type P2X7A subunits and alternatively spliced subunits (P2X7B, P2X7E, P2X7J, and P2X7L) that have been confirmed in humans. The study supports laboratory research by providing insight into the structure and flexibility of the heterotrimeric alternatively spliced receptors in a simulated environment and may thereby aid structure-guided drug design.

**Abstract:** P2X7 receptors (P2X7Rs) are membrane-bound ATP-gated ion channels that are composed of three subunits. Different subunit structures may be expressed due to alternative splicing of the *P2RX7* gene, altering the receptor’s function when combined with the wild-type P2X7A subunits. In this study, the application of the deep-learning method, AlphaFold2-Multimer (AF2M), for the generation of trimeric P2X7Rs was first validated by comparing an AF2M-generated rat wild-type P2X7A receptor with a structure determined by cryogenic electron microscopy (Protein Data Bank Identification: 6U9V). The results suggested AF2M could firstly, accurately predict the structures of P2X7Rs and secondly, accurately identify the highest quality model through the ranking system. Subsequently, AF2M was used to generate models of heterotrimeric alternatively spliced P2X7Rs consisting of one or two wild-type P2X7A subunits in combination with one or two P2X7B, P2X7E, P2X7J, and P2X7L splice variant subunits. The top-ranking models were deemed valid based on AF2M’s confidence measures, stability in molecular dynamics simulations, and consistent flexibility of the conserved regions between the models. Visual analysis of the heterotrimeric receptors identified missing residues in the ATP binding sites of the P2X7E, P2X7J, and P2X7L splice variants, likely translating into dysfunctional binding sites. Overall, the models produced in this study (available as supplementary material) unlock the possibility of structure-based studies into the heterotrimeric P2X7Rs.

## Introduction

P2X7 receptors (P2X7Rs) have been associated with cancers such as leukaemia (Zhang, et al., 2004), autoimmune conditions such as rheumatoid arthritis (Bahari, et al., 2021), trauma (Liu, et al., 2017), and neurodegenerative conditions such as Alzheimer’s disease (Martin, et al., 2019). The receptors are a subtype of the P2X receptors; a class of membrane-bound ATP-gated ion channels that are composed of three subunits (Fig. 1) (Fryatt, et al., 2019). Each P2X7R contains three ATP and three allosteric binding sites (Fig. 2) (McCarthy, et al., 2019). Activation by ATP results in the efflux of K^+^ and influx of Na^+^ and Ca^2+^ ions that correspondingly enhance cell proliferation and trigger a pro-inflammatory cascade (Pegoraro, et al., 2021). It is widely supported that prolonged ATP activation correlates with cell apoptosis however, the process through which this occurs is disputed. One theory suggests cell apoptosis occurs because the receptor’s channel diameter increases to form a pore that *in vitro* allows molecules up to 900 Da, such as YO-PRO-1 or ethidium bromide to pass through the cell membrane (Adinolfi, et al., 2005, Notomi, et al., 2011, Virginio, et al., 1999). Riedel, et al. (2007) disputes this pore formation theory, proposing that single channel kinetics and permeation properties are independent of receptor activation. Li, et al. (2015), in turn, has suggested that cell apoptosis results from time-dependent alterations in intracellular ion concentrations due to shifts in equilibrium associated with prolonged P2X7R activation.

**Fig. 1.**
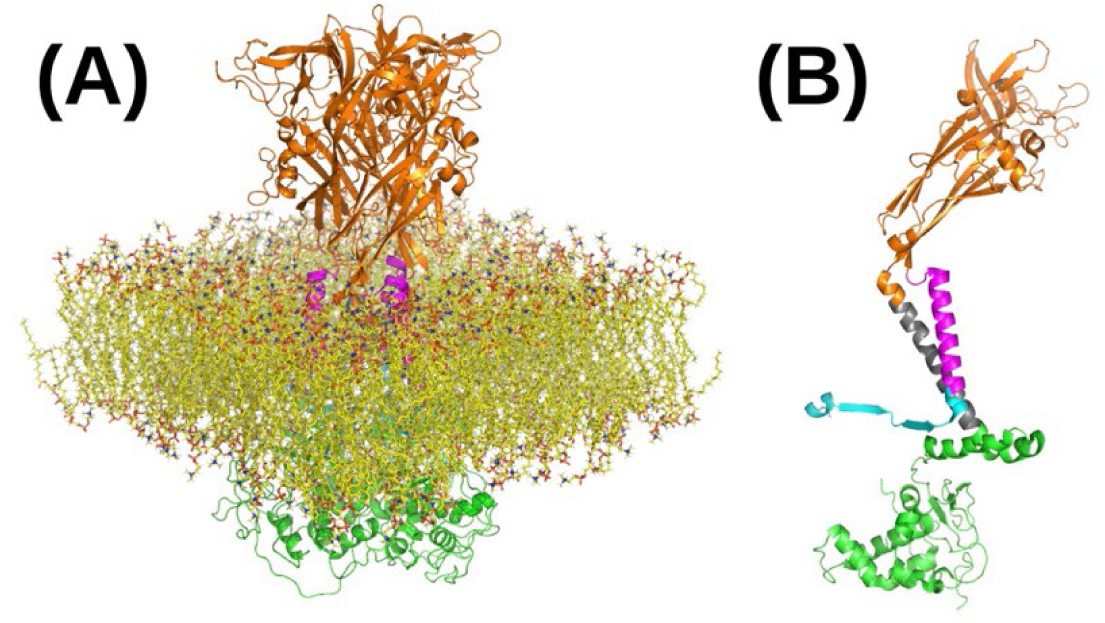
Apo closed state rat wild-type P2X7R structure determined by cryogenic electron microscopy (Protein Data Bank Identification: 6U9V) (McCarthy, et al., 2019) showing the amino terminal domain (cyan), transmembrane domain 1 (magenta), extracellular domain (orange), transmembrane domain 2 (grey), and carboxy terminal domain (green). **(A)** The P2X7R in a cell membrane (yellow). **(B)** An individual P2X7R subunit. Figures were generated using the PyMOL molecular graphics system, version 2.5.3, Schrödinger, LLC.

**Fig. 2.**
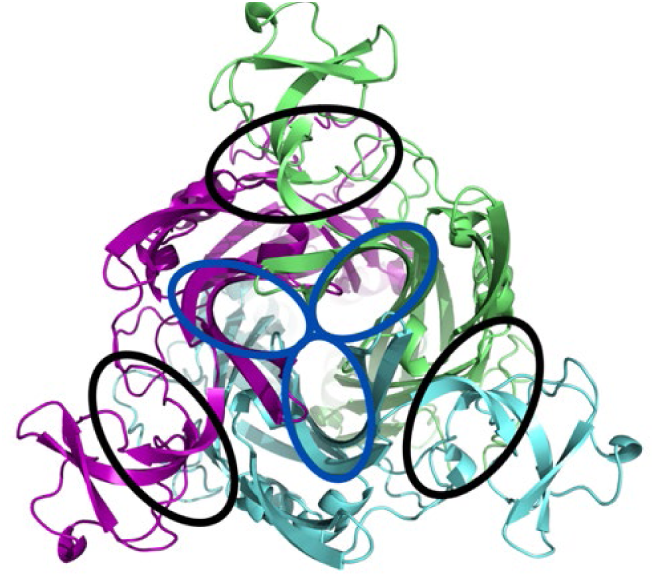
Birdseye view of the open state rat P2X7R determined by cryogenic electron microscopy (Protein Data Bank Identification: 6U9W). The three occupied ATP binding sites are circled in black. Each ATP binding site is located between two subunits (involving Lys64 and Lys66 in one subunit, and Asn292, Arg294 and Lys311 in the adjacent subunit) (McCarthy, et al., 2019). The three empty allosteric binding sites are circled in dark blue and are located within each subunit (involving Phe95, Phe103, Met105, Phe293, and Val312) (Illes, et al., 2021, Keystone, et al., 2012, Stock, et al., 2012). Chains A, B, and C are coloured green, cyan, and magenta. The figure was generated using the PyMOL molecular graphics system, version 2.5.3, Schrödinger, LLC.

The P2X7R structure is complex because of alternative splicing of the *P2RX7* gene that results in the expression of different subunit structures. Knowledge of the different P2X7R structures is key to supporting laboratory research and improving our understanding of the receptor’s role in disease. Five different subunits are currently known to translate from the *P2RX7* gene (Table 1; Fig. 3) (Cheewatrakoolpong, et al., 2005, Feng, et al., 2006, Skarratt, et al., 2020). The primary structure of the wild-type P2X7A subunit consists of an intracellular amino terminal domain (ATD), transmembrane domain (TMD) 1, extracellular domain (ECD), TMD 2, and intracellular carboxy terminal domain (CTD) (Fig. 1; Fig. 3) (Cheewatrakoolpong, et al., 2005, Adinolfi, et al., 2010). Comparatively, P2X7B subunits lack the CTD (Cheewatrakoolpong, et al., 2005); P2X7E subunits are missing part of the ECD and CTD (Skarratt, et al., 2020); P2X7J subunits are missing part of the ECD, the entire TMD 2, and CTD (Feng, et al., 2006); and P2X7L subunits are missing part of the ECD (Skarratt, et al., 2020).

**Fig. 3.**
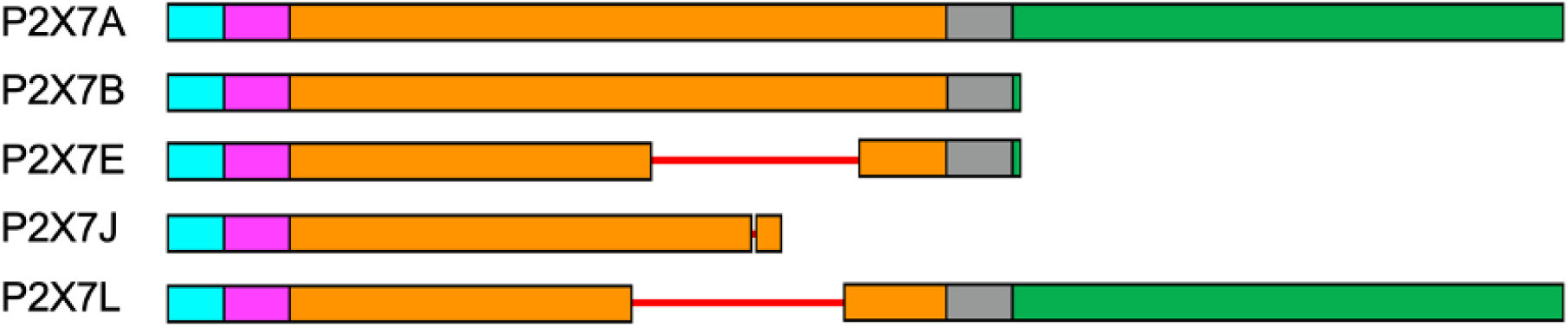
Schematic representation of the wild-type P2X7A (UniProtKB: Q99572-1), P2X7B (UniProtKB: Q99572-2), P2X7E (UniProtKB: Q99572-5), P2X7J (GenBank: DQ399293.1), and P2X7L (GenBank: MK465687.1) subunit amino acid sequences generated using NCBI Multiple Sequence Alignment Viewer, version 1.22.0, National Center for Biotechnology Information, Rockville, MD, USA (Papadopoulos and Agarwala, 2007). Residues 1 to 26 fold to form the intracellular ATD (blue), 27 to 51 the TMD 1 (pink), 52 to 330 the extracellular domain (orange), 331 to 355 the TMD 2 (grey), and 356 to 595 the intracellular CTD (green) (Adinolfi, et al., 2010).

**Table 1.** The five different subunits translated from the *P2RX7* gene in humans.

| Splice variants | Sequence length | Accession codes |
| --- | --- | --- |
| Wild-type P2X7A | 595 | UniprotKB: Q99572-1<sup>a</sup> |
| P2X7B | 364 | UniprotKB: Q99572-2<sup>b</sup> |
| P2X7E | 275 | UniprotKB: Q99572-5<sup>b</sup> |
| P2X7J | 258 | GenBank: DQ399293.1<sup>c</sup> |
| P2X7L | 506 | GenBank: MK465687.1<sup>d</sup> |

<sup>a</sup> (Rassendren, et al., 1997); <sup>b</sup> (Cheewatrakoolpong, et al., 2005); <sup>c</sup> (Feng, et al., 2006);
<sup>d</sup> (Skarratt, et al., 2020)

Current research suggests P2X7B, P2X7E, P2X7J, and P2X7L subunits can associate with wild-type P2X7A subunits to form heterotrimeric receptors (P2X7A/A/B, P2X7A/B/B, P2X7A/A/E, P2X7A/E/E, P2X7A/A/J, P2X7A/J/J, P2X7A/A/L, and P2X7A/L/L receptors) (Adinolfi, et al., 2010, Skarratt, et al., 2020, Feng, et al., 2006). ATP is more potent to P2X7B heterotrimers than wild-type P2X7A homotrimers (Adinolfi, et al., 2005). The P2X7E heterotrimers maintain the same level of function, the P2X7J heterotrimers are non-functional, and the P2X7L heterotrimers maintain the ion channel function but have reduced pore formation capabilities in comparison to the wild-type P2X7A homotrimers (Skarratt, et al., 2020, Feng, et al., 2006).

P2X7B expression has been widely studied in cancer (Pegoraro, et al., 2021). For example, a study observing P2X7R expression in myelodysplastic syndrome and acute myeloid leukaemia reported significantly higher wild-type P2X7A and P2X7B mRNA expression in acute myeloid leukaemia patients compared to those with myelodysplastic syndrome (Pegoraro, et al., 2020). Furthermore, treatment-unresponsive patients were found to have lower levels of the wild-type P2X7A mRNA and higher P2X7B mRNA expression, bringing to question whether a P2X7B-selective antagonist could be beneficial in combination with current treatment regimens (Pegoraro, et al., 2020). P2X7B has also been associated with cell proliferation in osteosarcoma (Giuliani, et al., 2014) and neuroblastoma (Ulrich, et al., 2018) making it an interesting focus for further research. P2X7J, despite its non-functionality, has been proposed as a potential biomarker for cervical cancer (Feng, et al., 2006). P2X7E and P2X7L currently have limited published information relating to disease associations but warrant further investigation based on their ability to express as functional proteins in combination with wild-type P2X7A subunits. For further information on P2X7R disease associations refer to De Salis, et al. (2022).

Despite studies showing that the P2X7R plays a critical role in human disease, results of P2X7R antagonists in human trials have been disappointing. This is likely to have resulted from drug-target interaction studies that used the wild-type P2X7A receptor and ignored the alternatively spliced isoforms. The structures of P2X7R antagonists are diverse, indicating that in addition to the ATP binding site and an inhibitory drug-binding pocket (Karasawa and Kawate, 2016), there are multiple other unmapped inhibitor binding sites. There is an opportunity to identify drug targets informed by the effects of alternative splicing considering the enormous promise of inhibiting P2X7Rs in human diseases. An exciting new deep learning-based modelling method, AlphaFold2-Multimer (AF2M) (Evans, et al., 2022, Jumper, et al., 2021, Mirdita, et al., 2022), has provided a pathway to elucidate these alternatively spliced structures. AF2M overcomes some of the limitations of traditional homology modelling which require high similarity templates to inform model generation (Hilbert, et al., 1993). In turn, the models may be used to aid the determination of experimental structures by acting as templates for molecular replacement in X-ray crystallography, and backbone tracing in cryogenic electron microscopy (cryo-EM) (Kryshtafovych, et al., 2021).

This study aims to validate the use of AF2M as a tool for P2X7R structure prediction, and then determine the structure of the alternatively spliced heterotrimeric receptors that express in humans (P2X7A/(A/B)/B, P2X7A/(A/E)/E, P2X7A/(A/J)/J, and P2X7A/(A/L)/L receptors) to complement current *in vitro* research into the heterotrimeric receptors. This study hypothesises that the unique structures of the heterotrimeric receptors can be modelled using AF2M and validated using AF2M’s confidence measures and molecular dynamics (MD) simulations.

## Materials and methods

The most recently published and highest quality experimentally obtained P2X7R structure is the apo closed state rat P2X7R determined by cryo-EM (Protein Data Bank Identification (PDB ID): 6U9V) (McCarthy, et al., 2019). Therefore, this structure was used as the experimental control for validation of the modelling approach. The residue and secondary structure numbering were assigned according to the wild-type P2X7A subunits. Refer to Table 2 for definitions on the validation criteria used and their cut-offs.

**Table 2.**
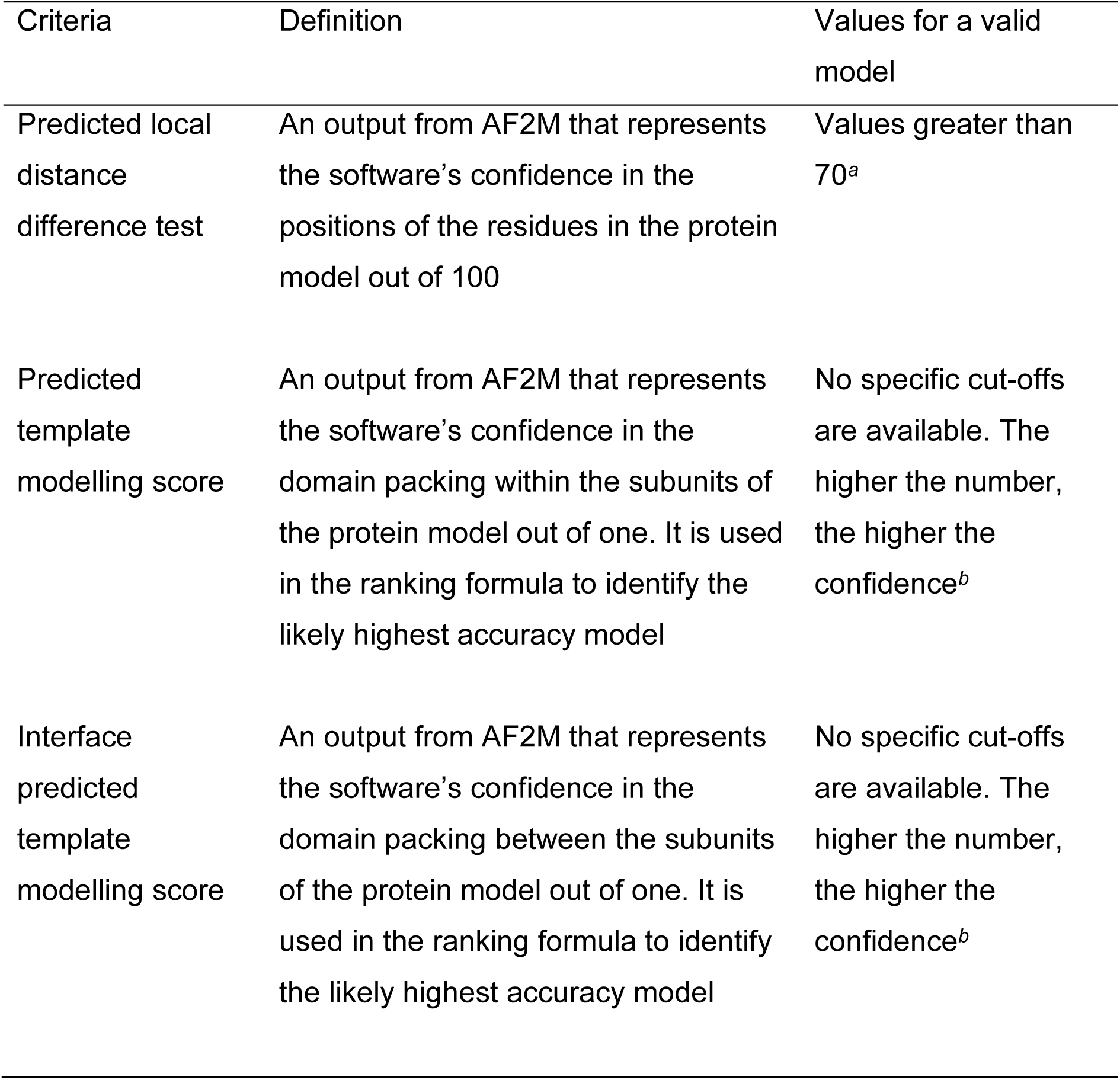
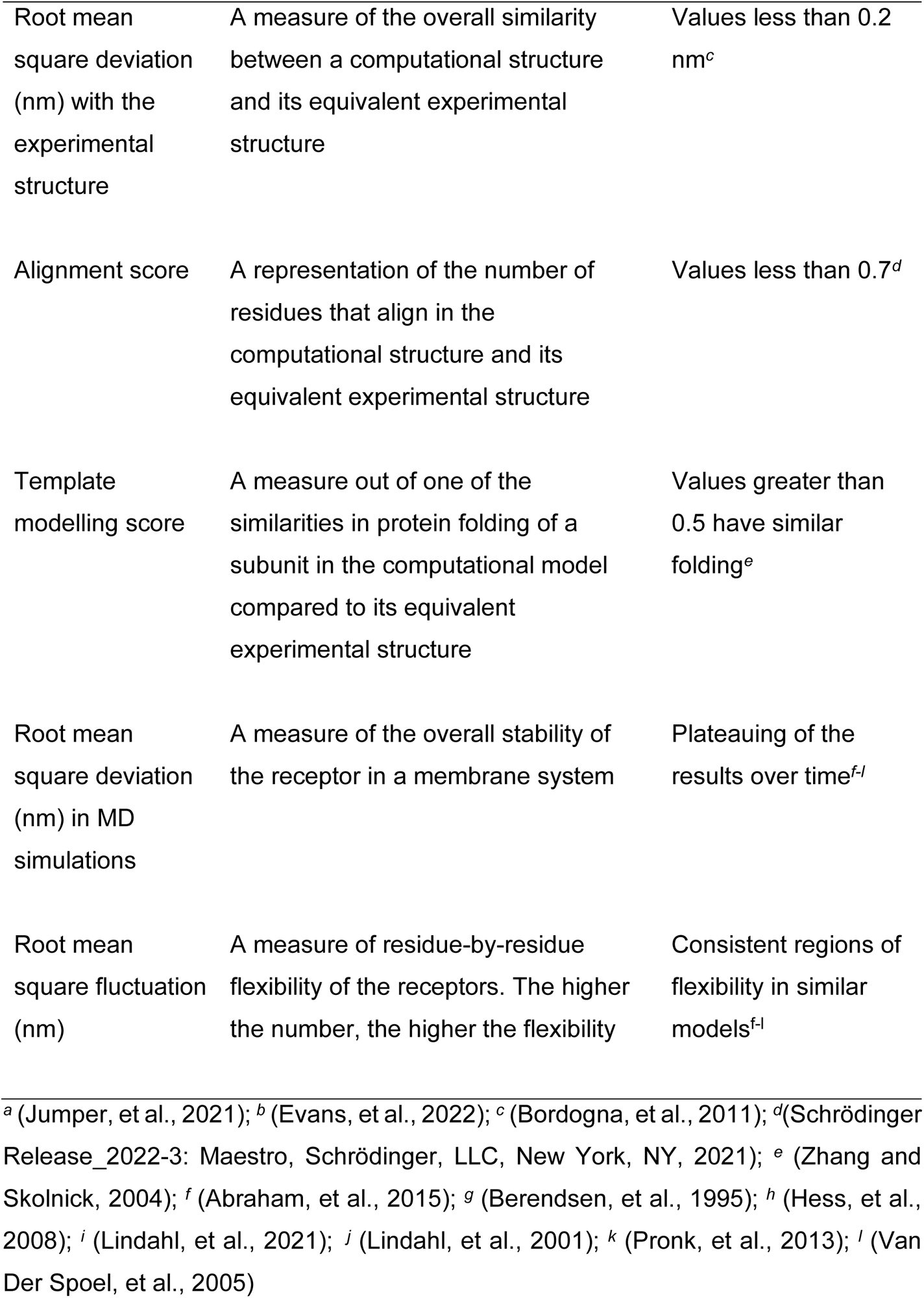
Definitions of the validation criteria used for the modelled structures.

### Method validation

AF2M was accessed as ColabFold 1.3.0 on the Australian National Computational Infrastructure Gadi supercomputer, using NVIDIA V100 graphics processing unit nodes (*gpuvolta* nodes) with 12 Intel Xeon Cascade Lake cores and 64GB of random-access memory per graphics processing unit (Evans, et al., 2022, Jumper, et al., 2021, Mirdita, et al., 2022). The rat wild-type P2X7A subunit amino acid sequence (uniprotKB: Q64663) was retrieved from UniProt (The UniProt, 2021) and submitted as a full receptor (three wild-type P2X7A amino acid sequences were entered) into AF2M. The structure generation process was informed by completing three paired and unpaired multiple sequence alignments (MSA) on each of the three entered amino acid sequences, searching the UniRef100 and environmental sequences databases using Many-against-Many sequence searching (Mirdita, et al., 2021, Mirdita, et al., 2019, Steinegger and Söding, 2018, Steinegger and Söding, 2017). Three .*a3m* MSA files were generated, one for each of the amino acid sequences. The files were manually combined into a single .*a3m* file for input into AF2M. 48 Evoformer blocks were used to process the MSA information with three ensemble iterations to produce final MSA and pair representations that were fed into eight structure module blocks to predict five protein structures and output a per-residue confidence score (the predicted local distance difference test (pLDDT)) (Table 3). The models were recycled through twenty iterative refinement cycles and relaxed using an AMBER protocol (Case, et al., 2005, Salomon-Ferrer, et al., 2013) to improve the likelihood of obtaining accurate structures and to address potential AF2M issues by relaxing side chain conformations (Jumper, et al., 2021, Evans, et al., 2022). The five models were ranked according to formula 1 (Evans, et al., 2022).

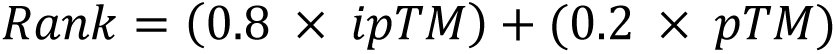

**Table 3.** Interpretation of the pLDDT score generated by AF2M for each residue in the models (Jumper, et al., 2021)

| pLDDT score | AF2M confidence |
| --- | --- |
| 90-100 | Very high confidence |
| 70-89 | High confidence |
| 50-69 | Low confidence |
| 0-49 | Very low confidence |

**Formula 1** The formula used to calculate the rank of the AF2M models. The predicted template modelling (pTM) scores and the interface pTM (ipTM) scores are measures of AF2M’s confidence in the domain packing within and between the receptor subunits respectively with scores ranging from zero (low confidence) to one (high confidence) (Evans, et al., 2022).

The root mean square deviation (RMSD_experimental_) between the five ranked AF2M rat wild-type P2X7A receptor models and the experimental control were calculated using formula 2 via the Protein Structure Alignment function in Maestro (Schrödinger Release_2022-3: Maestro, Schrödinger, LLC, New York, NY, 2021) to measure their overall similarity.

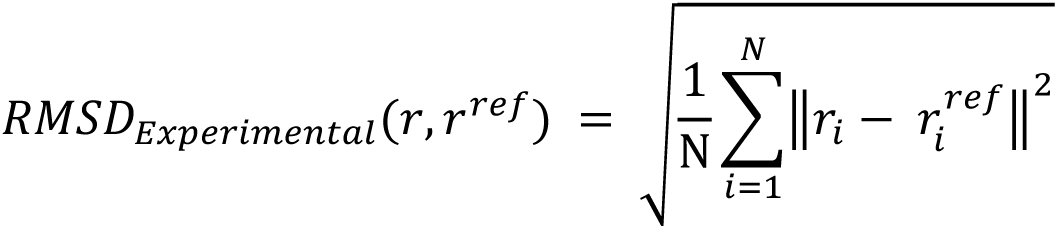

**Formula 2** Root mean square deviation between a computational model and its experimental structure (RMSD_experimental_) where N is the number of aligned residues, and r_i_ – r_i_^ref^ is the difference in the positions of the residue *i* in the AF2M models (r) compared to the experimental control (r^ref^) (Schrödinger Release_2022-3: Maestro, Schrödinger, LLC, New York, NY, 2021).

The template modelling (TM) scores of the AF2M models compared to the experimental control were calculated using formula 3 to measure the similarities in subunit folding (Xu and Zhang, 2010, Zhang and Skolnick, 2004).

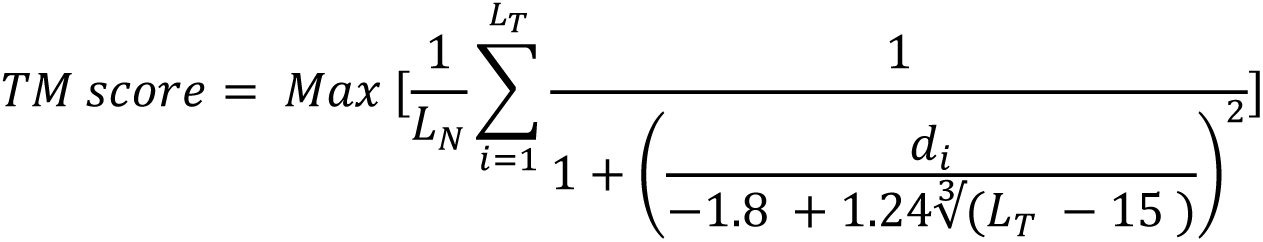

**Formula 3** The template modelling (TM) score between a computational model and its experimental structure where L_N_ is the length of the primary sequence of the AF2M subunits, L_T_ is the number of residues that correspond with the experimental control’s subunits, d_i_ is the distance between residue i in the AF2M models and its equivalent residue in the experimental control, and max represents the maximum d_i_ (Xu and Zhang, 2010, Zhang and Skolnick, 2004).

ColabFold 1.3.0 does not have a native function to search for structural templates to inform the structure generation process. Thus, a custom template database was developed (containing the experimental structures of the apo closed state rat P2X7R (PDB ID: 6U9V) (McCarthy, et al., 2019), giant panda P2X7R (PDB ID: 5U1L) (Karasawa and Kawate, 2016), and zebrafish P2X4 receptor (PDB ID: 3I5D) (Kawate, et al., 2009)) and tested to determine the effect of using templates on the quality of the generated models. The RMSD_experimental_ of the top-ranked models generated with and without templates were subsequently compared.

### Heterotrimeric P2X7R models

The human wild-type P2X7A (Rassendren, et al., 1997), P2X7B (Cheewatrakoolpong, et al., 2005), P2X7E (Cheewatrakoolpong, et al., 2005), P2X7J (Feng, et al., 2006), and P2X7L (Skarratt, et al., 2020) amino acid sequences were downloaded from the UniProt (The UniProt, 2021) and GenBank (Clark, et al., 2016) databases. The method described above to model the rat P2X7R (with no custom template) was replicated to model the human wild-type P2X7A, P2X7A/(A/B)/B, P2X7A/(A/E)/E, P2X7A/(A/J)/J, and P2X7A/(A/L)/L receptors, altering the entered amino acid sequences to match the respective subunits.

The top-ranked models for each of the receptors were selected to study further. The structures of the heterotrimeric receptor models were compared to the human wild-type P2X7A receptor model and their pLDDT scores were analysed.

### P2X7R molecular dynamics simulations

Receptor preparation was carried out using the Protein Preparation Wizard in Maestro (Schrödinger Release_2022-3: Protein Preparation Wizard; Epik, Schrödinger, LLC, New York, NY, 2021; Impact, Schrödinger, LLC, New York, NY; Prime, Schrödinger, LLC, New York, NY, 2021) (Sastry, et al., 2013). The experimental control, human wild-type P2X7A, and heterotrimeric P2X7R models were pre-processed: assigned bond orders, aligned with the corresponding P2X7R structures in the Orientations of Proteins in Membranes (OPM) database (Lomize, et al., 2012, Olsson, et al., 2011), and structural issues such as missing side chains and missing hydrogen atoms were corrected. Hydrogen bonds were created and optimised using PROPKA at pH 7.4 (Søndergaard, et al., 2011). The heavy atoms and hydrogens were minimised using the OPLS4 force field (Lu, et al., 2021) with a convergence threshold offset to 0.03 nm to address clashes that may have occurred from the addition of the side chains and hydrogens.

The experimental control is missing loops from residues 76 to 80, and 443 to 469 (McCarthy, et al., 2019). These were filled in automatically using Maestro Crosslink Proteins, predicting the loop conformation using simple *de novo* loop creation with an implicit solvent model in the energy calculation (Schrödinger Release_2022-3: Maestro, Schrödinger, LLC, New York, NY, 2021). The control was prepared again using Maestro’s Protein Preparation Wizard to address atom overlaps in the regions with added loops (Schrödinger Release_2022-3: Protein Preparation Wizard; Epik, Schrödinger, LLC, New York, NY, 2021; Impact, Schrödinger, LLC, New York, NY; Prime, Schrödinger, LLC, New York, NY, 2021) (Sastry, et al., 2013).

The receptor-membrane systems were prepared using a CHARMM36m force field in CHARMM-GUI (Brooks, et al., 2009, Jo, et al., 2007, Jo, et al., 2008, Jo, et al., 2009, Lee, et al., 2016, Lee, et al., 2019, Wu, et al., 2014). Residues 360, 362, 363, 371, 373, 374, and 377 in the experimental control (McCarthy, et al., 2019); residues 4, 360, 362, 363, 374, and 377 in the human wild-type P2X7A and P2X7L subunits; residues 4, 360, 362, and 363 in the P2X7B and P2X7E subunits; and residue 4 in the P2X7J subunits were palmitoylated.

The receptors were inserted into lipid bilayers composed of 30% cholesterol (Ingólfsson, et al., 2014, Wilson, et al., 2021) (16% upper leaflet, 84 lipids; 14% lower leaflet, 70 lipids) and 70% 1-palmitoyl-2-oleoyl-sn-glycero-3-phosphocholine (POPC) (36% upper leaflet, 189 lipids; 34% lower leaflet, 170 lipids). The receptors and membranes were enclosed in rectangular systems (approximately 13 x 13 x 22 nm) with a water thickness above and below the receptor of 22.5 nm. Na^+^ and Cl^-^ ions were added to a concentration of 150 mM via random replacement of water molecules. Additional Na^+^ or Cl^-^ ions were added until the system charge was neutralised.

MD simulations were performed in GROMACS 2021.4 on NCI Gadi *gpuvolta* nodes using the CHARMM-GUI prepared systems to determine the stability of the AF2M-generated models in a POPC and cholesterol membrane system (Abraham, et al., 2015, Berendsen, et al., 1995, Hess, et al., 2008, Lindahl, et al., 2021, Lindahl, et al., 2001, Pronk, et al., 2013, Van Der Spoel, et al., 2005). Minimisation, equilibration, and production steps were conducted using the same interaction and constraint algorithms as follows: particle neighbour lists with a neighbour search cut-off of 1.2 nm were updated every twenty steps during minimisation and every ten steps during equilibration and production; the particle pair interactions for each list were calculated using the Verlet cut-off scheme (Páll and Hess, 2013); hydrogen bonds were defined as constraints using the LINCS algorithm (Hess, et al., 1997); and the interactions were constrained with a neighbour search cut-off of 1.2 nm using the Force-switch modifier (Abraham, et al., 2015, Berendsen, et al., 1995, Hess, et al., 2008, Lindahl, et al., 2021, Lindahl, et al., 2001, Pronk, et al., 2013, Van Der Spoel, et al., 2005) for van der Waal interactions and the fast smooth Particle-Mesh Ewald method (Essmann, et al., 1995) for electrostatic interactions.

Minimisation of the receptor-membrane systems adjusted the atomic arrangement to one that was energetically favourable (maximum force < 1000 kJ mol^-1^ nm^-1^) using the steepest descents method (Cauchy, 1847) with a maximum of 5000 steps. Six cycles of equilibration were performed to remove the centre of mass motion and equilibrate the receptor-membrane systems environmental factors using a time step of 2 fs. A Berendsen thermostat (Berendsen, et al., 1984) was used to equilibrate the system at a constant temperature of 303.15 K with a coupling time constant of 1000 fs. A Berendsen barostat (Berendsen, et al., 1984) was used to equilibrate the receptor-membrane system at a constant pressure of 1 bar and compressibility of 4.50e^-5^ bar^-1^ with semi-isotropic pressure coupling and a time constant of 1000 fs. Three randomly seeded production simulations were conducted for each of the systems using a time step of 2 fs and a total of 125000000 steps for a total simulation time of 250 ns. A Nosé-Hoover thermostat (Hoover, 1985, Nosé, 1984) was used to maintain a constant temperature of 303.15 K with a coupling temperature fluctuation period of 1000 fs. A Parrinello-Rahman barostat (Parrinello and Rahman, 1981) was used to maintain a constant pressure of 1 bar and compressibility of 4.5e^-5^ bar^-1^ with a semi-isotropic pressure coupling and a temperature fluctuation period of 5000 fs.

The simulation trajectories were post-processed using *gmx trjconv* in GROMACS 2021.4 (Abraham, et al., 2015, Berendsen, et al., 1995, Hess, et al., 2008, Lindahl, et al., 2021, Lindahl, et al., 2001, Pronk, et al., 2013, Van Der Spoel, et al., 2005) to centre the protein in the membrane system. The RMSD of the 1250 receptor frames (every 0.2 ns of the 250 ns production simulation) compared to the initial receptor frame (t = 0 ns) (RMSD_MD_) for each of the protein-membrane systems were calculated via *gmx rms* on GROMACS 2021.4 using formula 4.

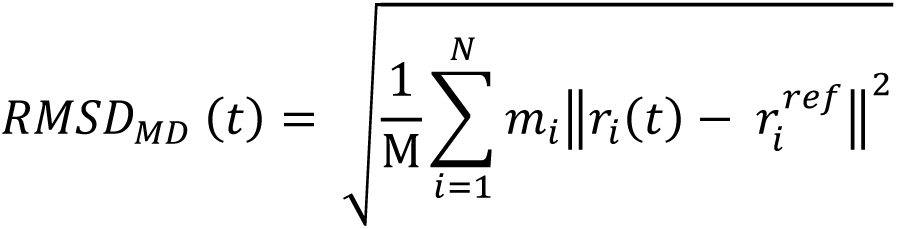

**Formula 4** The root mean square deviation across a MD simulation (RMSD_MD_) where M is the sum of the atomic masses, N is the number of atoms, m_i_ is the mass of atom i and r_i_(t) is the position of atom i at time t in comparison to the position of atom i in the initial receptor conformation r^ref^ (Abraham, et al., 2015, Berendsen, et al., 1995, Hess, et al., 2008, Lindahl, et al., 2021, Lindahl, et al., 2001, Pronk, et al., 2013, Van Der Spoel, et al., 2005).

The root mean square fluctuation (RMSF) was calculated for each residue in the receptors via *gmx rmsf* in GROMACS 2021.4 (Abraham, et al., 2015, Berendsen, et al., 1995, Hess, et al., 2008, Lindahl, et al., 2021, Lindahl, et al., 2001, Pronk, et al., 2013, Van Der Spoel, et al., 2005) from 150 to 250 ns (500 frames) in the production simulation using formula 5.

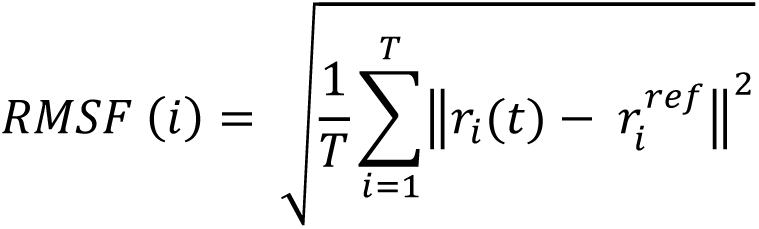

**Formula 5** The root mean square fluctuation (RMSF) across a MD simulation where T is the number of frames included in the calculation, r_i_(t) is the position of residue i at time t and r_i_^ref^ is the position of the residue in the initial receptor conformation (Abraham, et al., 2015, Berendsen, et al., 1995, Hess, et al., 2008, Lindahl, et al., 2021, Lindahl, et al., 2001, Pronk, et al., 2013, Van Der Spoel, et al., 2005).

The generated models are available in the supplementary material.

## Results

### Method validation

The method was validated by comparing the AF2M-generated rat P2X7R model with the equivalent model obtained via cryo-EM (experimental control). The top-ranked rat P2X7R model (ranked based on its pTM = 0.78 and ipTM = 0.77) was predicted with high per-residue confidence with a pLDDT value of 79.40. Alignment of the top-ranked model with the experimental control revealed that the two structures were similar as demonstrated by an RMSD_experimental_ value of 0.18 nm, alignment score of 0.15, and a TM score of 0.96 (Table 2).

Observation of the AF2M ranking method revealed that the method could accurately identify the model with the highest similarity to the experimental control as the top-ranking method using the pTM and ipTM scores. Of the five generated rat P2X7R models, the top-ranked model had an RMSD_experimental_ of 0.18 nm from the experimental control compared to 0.37 nm, 0.26 nm, 0.30 nm, and 0.32 nm for rank 2, 3, 4, and 5 respectively.

The addition of a custom template database to inform the structure generation process did not impact the quality of the AF2M rat P2X7R models. The difference in RMSD_experimental_ of the top-ranked models with and without templates was 0.01 nm.

### Heterotrimeric P2X7R models

The AF2M models of the top-ranked human wild-type P2X7A and heterotrimeric P2X7A/(A/B)/B, P2X7A/(A/E)/E, P2X7A/(A/J)/J, and P2X7A/(A/L)/L receptors are shown in Fig. 4. The elements of the wild-type P2X7A subunit secondary structure are outlined in Fig. 5 as a comparator for the splice variants. The primary and secondary structures of the subunits are outlined in Table 4.

**Fig. 4.**
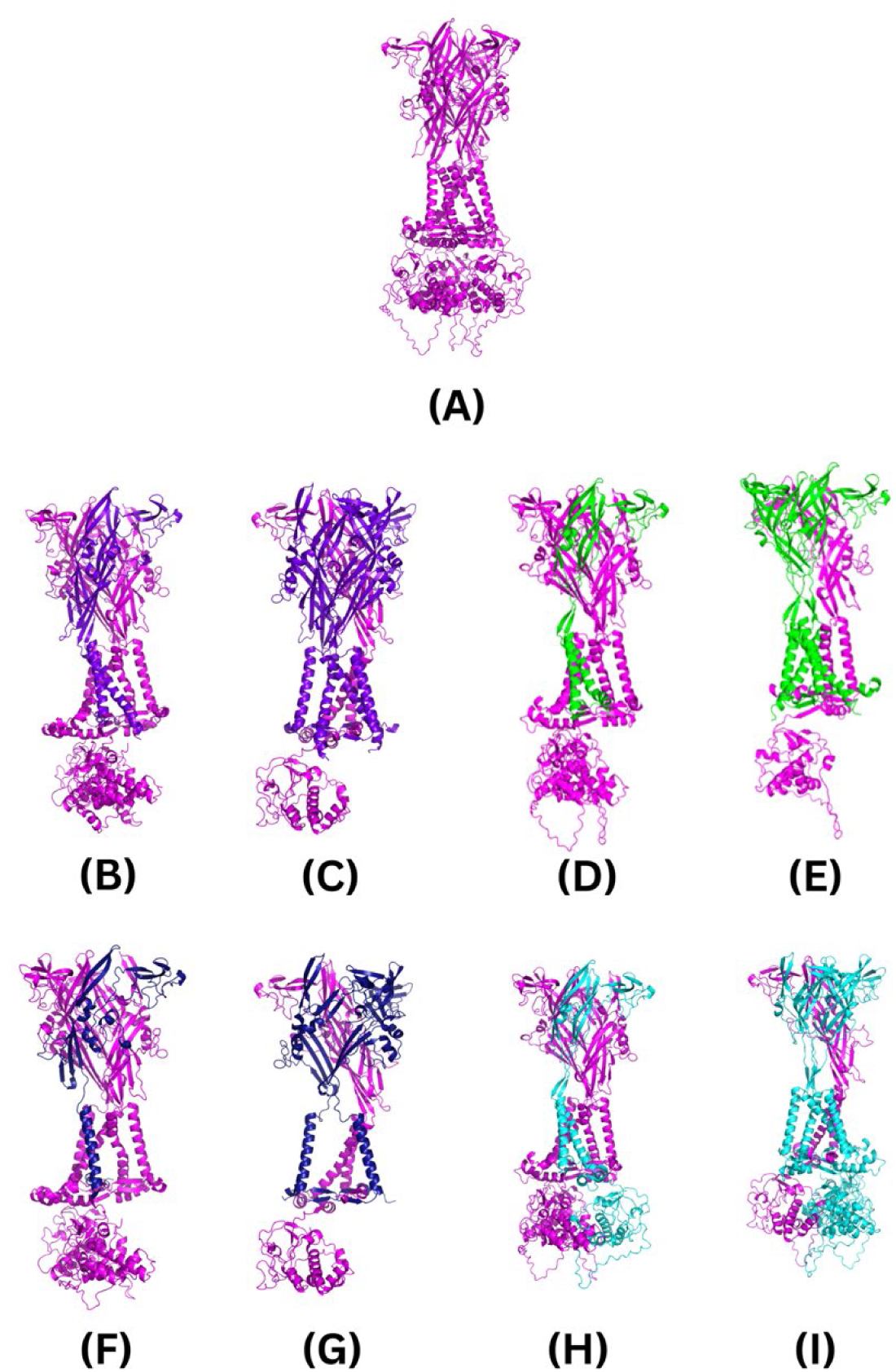
The AF2M-generated human P2X7R structures. The subunits are colour coded as follows: wild-type P2X7A is magenta; P2X7B is purple; P2X7E is green; P2X7J is dark blue; and P2X7L is cyan. **(A)** The wild-type P2X7A receptor; **(B)** P2X7A/A/B receptor; **(C)** P2X7A/B/B receptor; **(D)** P2X7A/A/E receptor; **(E)** P2X7A/E/E receptor; **(F)** P2X7A/A/J receptor; **(G)** P2X7A/J/J receptor; **(H)** P2X7A/A/L receptor; **(I)** and P2X7A/L/L receptor. Figures were generated using the PyMOL molecular graphics system, version 2.5.3, Schrödinger, LLC.

**Fig. 5.**
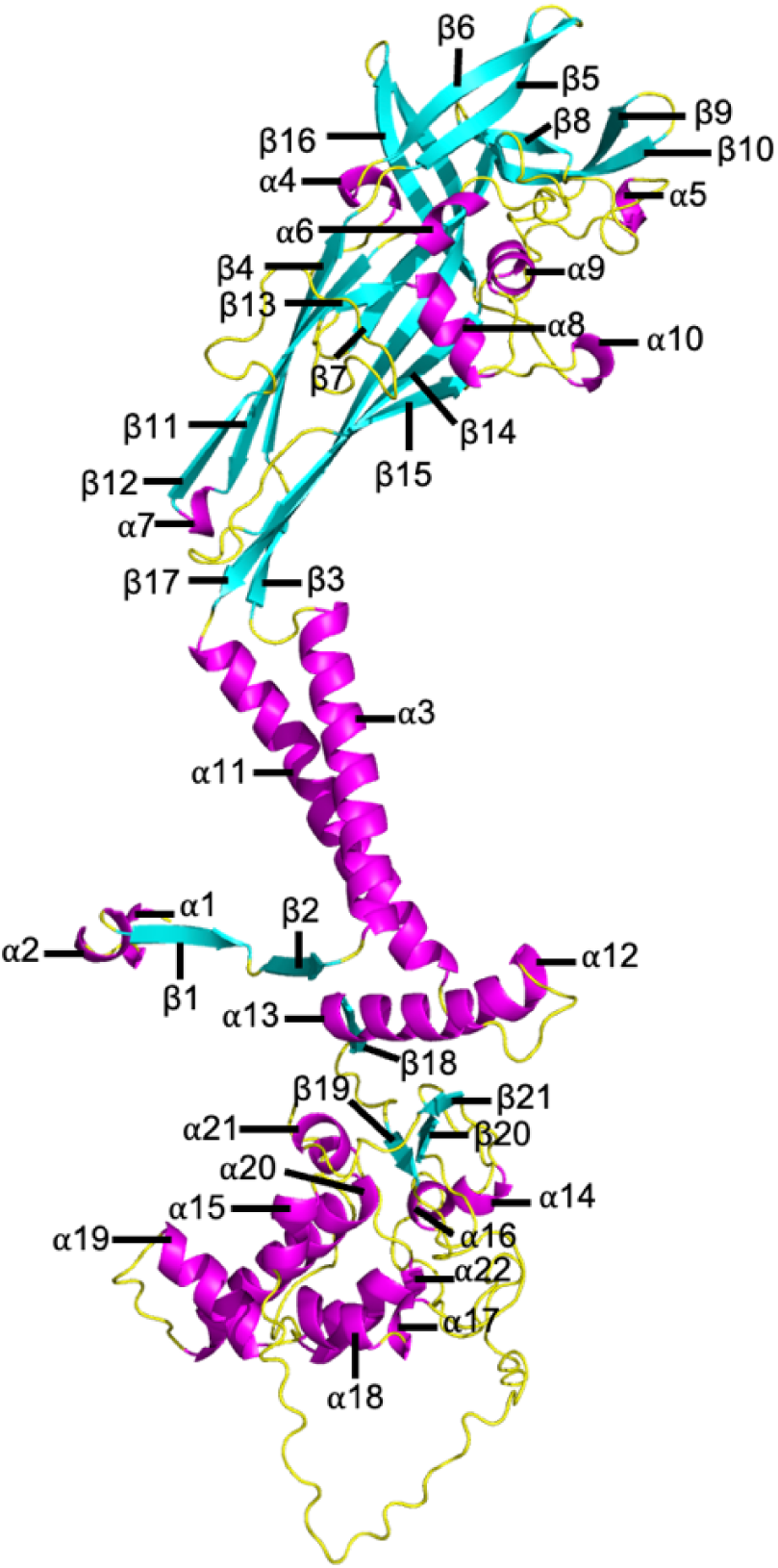
The AF2M-generated wild-type P2X7A subunit secondary structure with the naming of the individual segments adopted from McCarthy, et al. (2019). The figure was generated using the PyMOL molecular graphics system, version 2.5.3, Schrödinger, LLC.

**Table 4.** The primary and secondary structures of the P2X7R subunits. The missing regions of the splice variants are defined as areas which are absent in comparison to the wild-type P2X7A subunit.

| Splice variant | Primary structures | Secondary structures - $\alpha$ helices | Secondary structures - $\beta$ strands |
| --- | --- | --- | --- |
| Wild-type P2X7A | 595 amino acids | 22 $\alpha$ helices | 21 $\beta$ strands |
| P2X7B | 364 amino acids (missing 231 residues from residue 365 to 595) | 11 $\alpha$ helices (missing α12 to α22) | 17 $\beta$ strands (missing β18 to β21) |
| P2X7E | 275 amino acids (missing 320 residues from residue 206 to 294 and 365 to 595) | 7 $\alpha$ helices (missing α7 to α10, α12 to α22) | 16 $\beta$ strands (missing β12 to β15, β18 to β21, and β17 is split in two) |
| P2X7J | 258 amino acids (missing 337 residues from residue 259 to 595) | 10 $\alpha$ helices (missing α11 to α22) | 11 $\beta$ strands (missing β3, β7, β14 to β21) |
| P2X7L | 506 amino acids (missing 89 residues from residues 206 to 294) | 18 $\alpha$ helices (missing α7 to α10) | 18 $\beta$ strands (missing β12 to β15, and β17 is split in two) |

AF2M’s per-residue confidence score for each of the models was evaluated as a predicted measure of accuracy. All the top-ranked models were predicted with high per-residue confidence. The human P2X7A/B/B receptor was predicted with the highest confidence (pLDDT = 85.0), followed by the wild-type P2X7A (pLDDT = 83.5), P2X7A/J/J (pLDDT = 83.2), P2X7A/A/B (pLDDT = 81.8), P2X7A/A/J (pLDDT = 81.6), P2X7A/A/E (pLDDT = 81.0), P2X7A/A/L (pLDDT = 79.9), P2X7A/E/E (pLDDT = 76.1), and P2X7A/L/L (pLDDT = 76.1) receptors. Confidence in the model’s accuracy fluctuated on a per-residue basis as shown in Fig. 6. Lower per-residue confidence was observed in the loop regions of the receptors. For example, the loop region between β21 and α16 in the CTD (corresponding with residues 433 to 472) of the wild-type P2X7A and P2X7L subunits had a low per-residue confidence score (Fig. 6). Lower per-residue confidence was also observed in the loop regions near the spliced-out sites of the splice variants (residues 172 to 205, 295 to 303, and 349 to 365 in P2X7E; residues 249 to 258 in P2X7J; and residues 172 to 205, and 295 to 303 in the P2X7L splice variants) (Fig. 6).

**Fig. 6.**
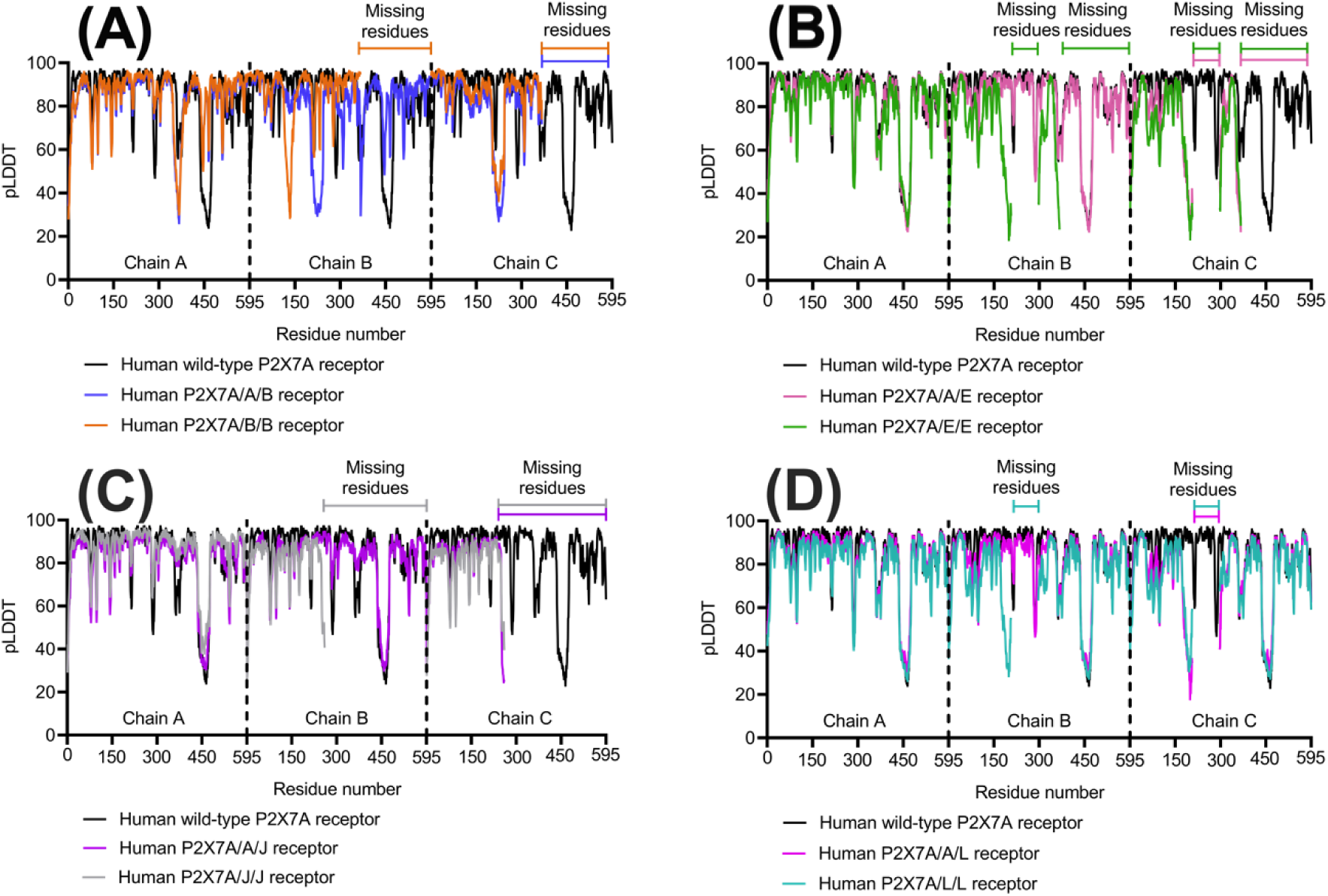
The wild-type P2X7A and heterotrimeric P2X7Rs pLDDT value per-residue out of 100 for each of the three subunits. The higher the pLDDT score, the higher the per-residue confidence. **(A)** The P2X7B heterotrimeric receptors compared to the wild-type P2X7A receptor. **(B)** The P2X7E heterotrimeric receptors compared to the wild-type P2X7A receptor. **(C)** The P2X7J heterotrimeric receptors compared to the wild-type P2X7A receptor. **(D)** The P2X7L heterotrimeric receptors compared to the wild-type P2X7A receptor. The graphs were generated using GraphPad Prism version 9.4.1 for MacOS, GraphPad Software, San Diego, California USA.

### P2X7R molecular dynamics simulations

MD simulation outputs were observed to identify the stability and flexibility of the receptors in a simulated membrane system. Similar stabilities were observed between the experimental control and the AF2M-generated human wild-type P2X7A and heterotrimeric receptors as shown in the RMSD_MD_ outputs. The RMSD_MD_ outputs increased within the first 1 ns of the simulations following the introduction of the initial particle velocities to the systems (Fig. 7). The outputs then plateaued to a mean *±* standard deviation from 150 to 250 ns in the simulation of 0.68 *±* 0.02 nm for the experimental control compared to 0.61 *±* 0.02 nm for the human wild-type P2X7A, 0.63 *±* 0.05 nm for the P2X7A/A/B, 0.66 *±* 0.04 nm for the P2X7A/A/L, 0.72 *±* 0.05 nm for the P2X7A/A/J, 0.76 *±* 0.02 nm for the P2X7A/A/E, 0.78 *±* 0.03 nm for the P2X7A/L/L, 0.79 *±* 0.06 nm for the P2X7A/B/B, 0.89 *±* 0.11 for the P2X7A/J/J, and 1.08 *±* 0.05 nm for the P2X7A/E/E receptors (Fig. 7).

**Fig. 7.**
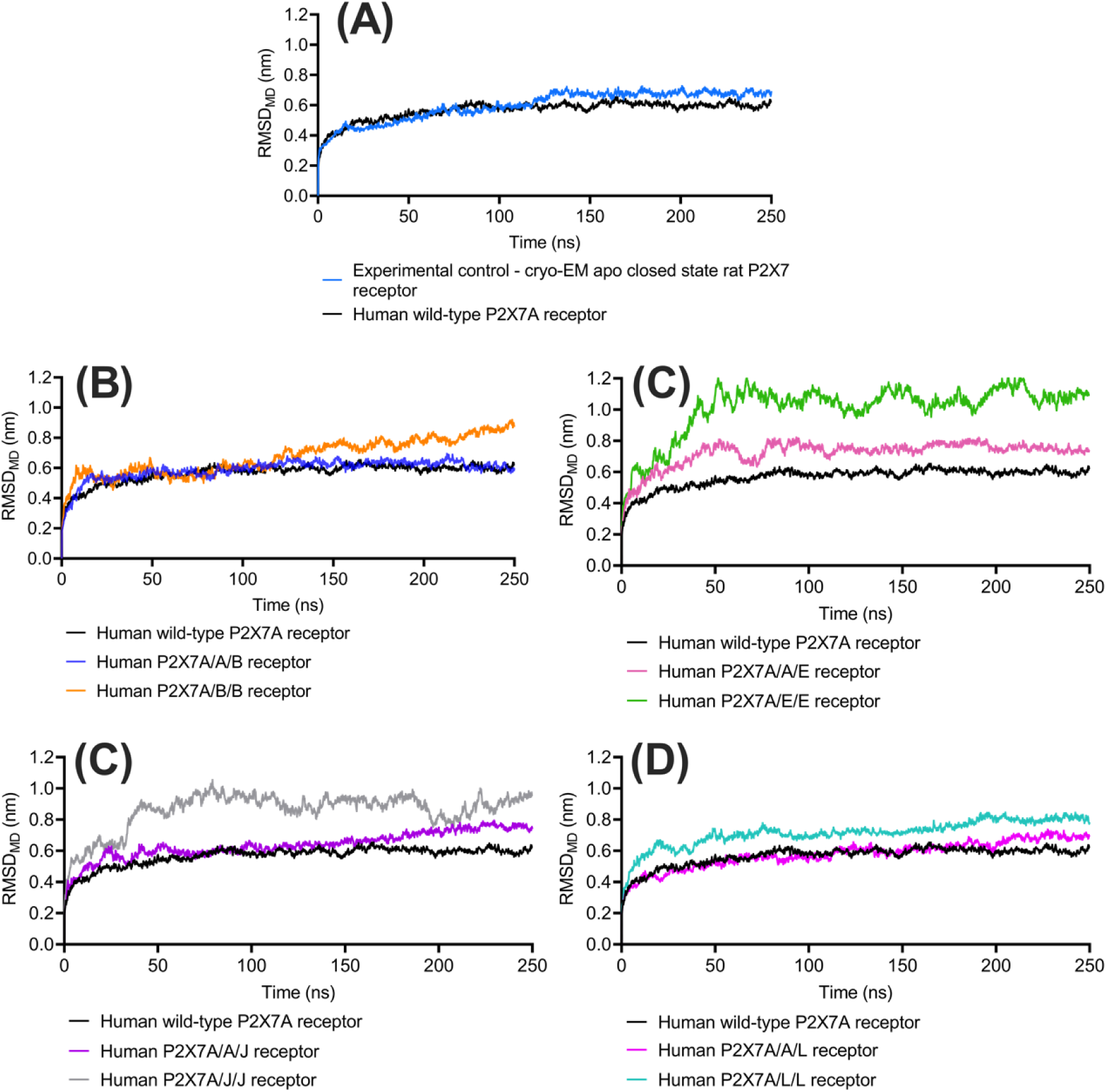
The wild-type P2X7A and heterotrimeric P2X7Rs average RMSD_MD_ (nm) over time (ns) in comparison to their initial receptor conformation. Plateauing of the results suggests the receptor is stabilising in the MD simulation trajectory. **(A)** The experimental control and human wild-type P2X7A receptor model. **(B)** The P2X7B heterotrimeric receptors compared to the wild-type P2X7A receptor. **(C)** The P2X7E heterotrimeric receptors compared to the wild-type P2X7A receptor. **(D)** The P2X7J heterotrimeric receptors compared to the wild-type P2X7A receptor. **(E)** The P2X7L heterotrimeric receptors compared to the wild-type P2X7A receptor. The graphs were generated using GraphPad Prism version 9.4.1 for MacOS, GraphPad Software, San Diego, California USA.

The flexibility of the residues in the receptors were consistent between the experimental control and the human wild-type P2X7A receptor AF2M-generated structure as shown in the RMSF outputs (Fig. 8A). A macroscopic inspection of the RMSF outputs for the experimental control and all the AF2M-generated structures identified peaks in flexibility in the loop segments of the receptors (Fig. 8). The highest RMSF peaks were observed in an unstructured region of the CTD in the wild-type P2X7A and P2X7L subunits from residues 446 to 492 (the loop between α9 and α10) that has not been elucidated in the rat P2X7R cryo-EM structure (Fig. 8E).

**Fig. 8.**
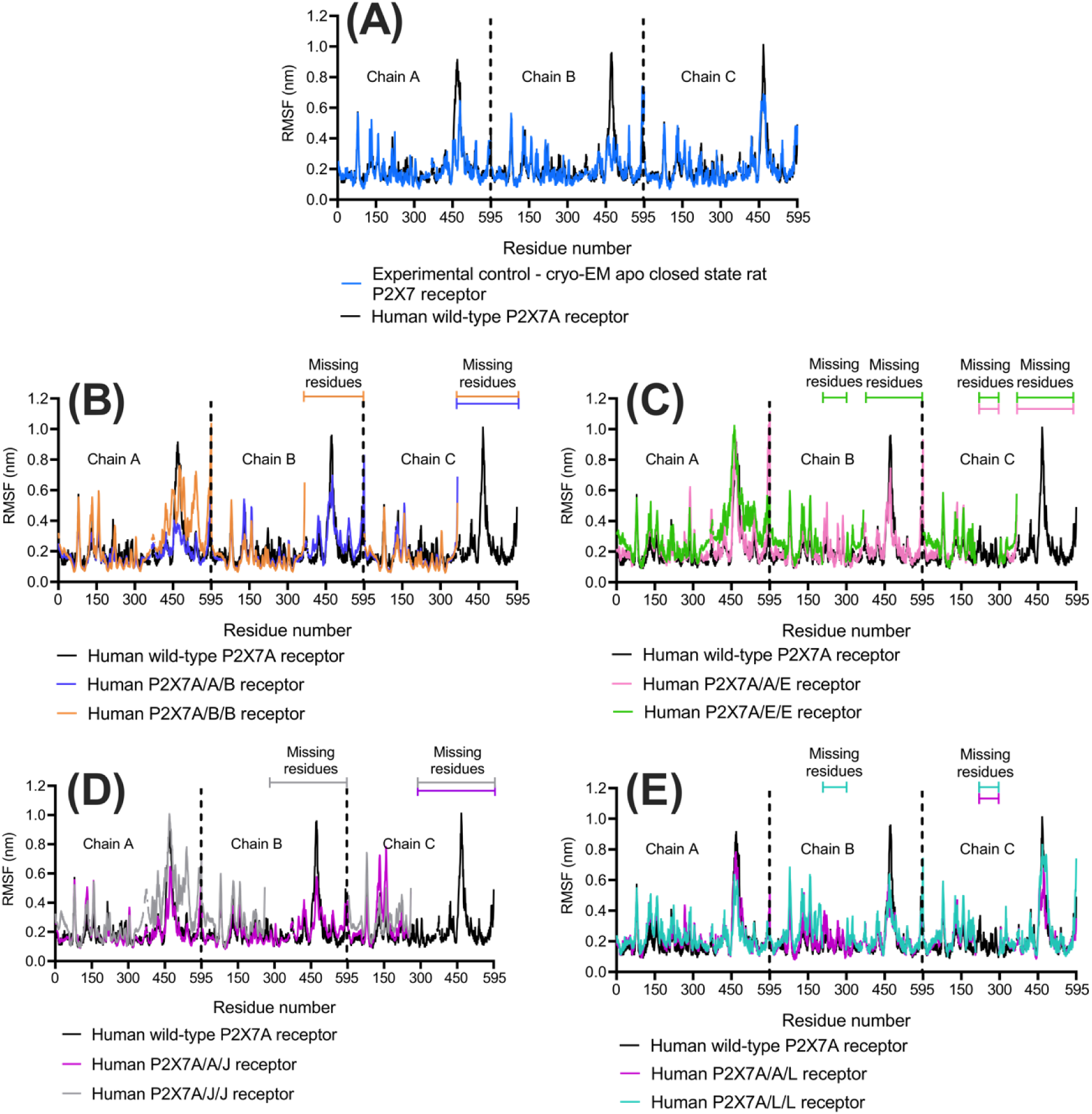
The wild-type and heterotrimeric P2X7 receptor’s RMSF (nm) per residue for each of the three subunits (chain A, B, C). Peaks represent regions of flexibility. The results were obtained from the MD simulations. **(A)** The experimental control and human wild-type P2X7A receptor model. **(B)** The P2X7B heterotrimeric receptors compared to the wild-type P2X7A receptor. **(C)** The P2X7E heterotrimeric receptors compared to the wild-type P2X7A receptor. **(D)** The P2X7J heterotrimeric receptors compared to the wild-type P2X7A receptor. **(E)** The P2X7L heterotrimeric receptors compared to the wild-type P2X7A receptor. The graphs were generated using GraphPad Prism version 9.4.1 for MacOS, GraphPad Software, San Diego, California USA.

The heterotrimeric receptors consisting of two splice variants were notably more flexible than those containing only one splice variant. In particular, the remaining intracellular regions of the P2X7A/B/B, P2X7A/E/E, and P2X7A/J/J heterotrimeric receptors were more flexible, consistent with these receptors being less restricted due to the absent large CTD that is present in the wild-type P2X7A and P2X7L subunits (Fig. 8). The splice variants also reported higher flexibilities in comparison to the wild-type P2X7A subunits in the residues located near the spliced-out sites (residues 358 to 364 in the P2X7B subunits; residues 117 to 203, 300 to 306 and 320 to 325 in the P2X7E subunits; residues 254 to 258 in the P2X7J subunits; and residues 117 to 203 and 300 to 306 in the P2X7L subunits) (Fig. 8).

### Model analysis

The P2X7E, P2X7J, and P2X7L splice variants with known modifications to the ATP binding region were further analysed. P2X7E and P2X7L are both missing the ATP binding regions from residues 206 to 294. P2X7J is missing all residues after residue 258. In the P2X7R models containing P2X7E (Fig. 9A), P2X7L (Fig. 9B), or P2X7J subunits (Fig. 9C), the ATP binding interfaces are disrupted. Different residues are located at the ATP binding region in each of the splice variants because of their altered structures (Table 5). The ATP binding sites in the P2X7E and P2X7L splice variants compared to the wild-type P2X7A subunits revealed that a key part of the ECD that forms the outer “lobe” of the ATP binding region is missing, while the inner two “lobes” of the binding region remain intact (Fig. 9C). P2X7J splice variants are missing the inner “lobes,” modifying the ATP binding site in a different manner. Accordingly, the P2X7A/A/E, P2X7A/A/L, and P2X7A/A/J receptors have one disrupted ATP binding site while the P2X7A/E/E, P2X7A/L/L, and P2X7A/J/J receptors have two disrupted ATP binding sites.

**Figure 9.**
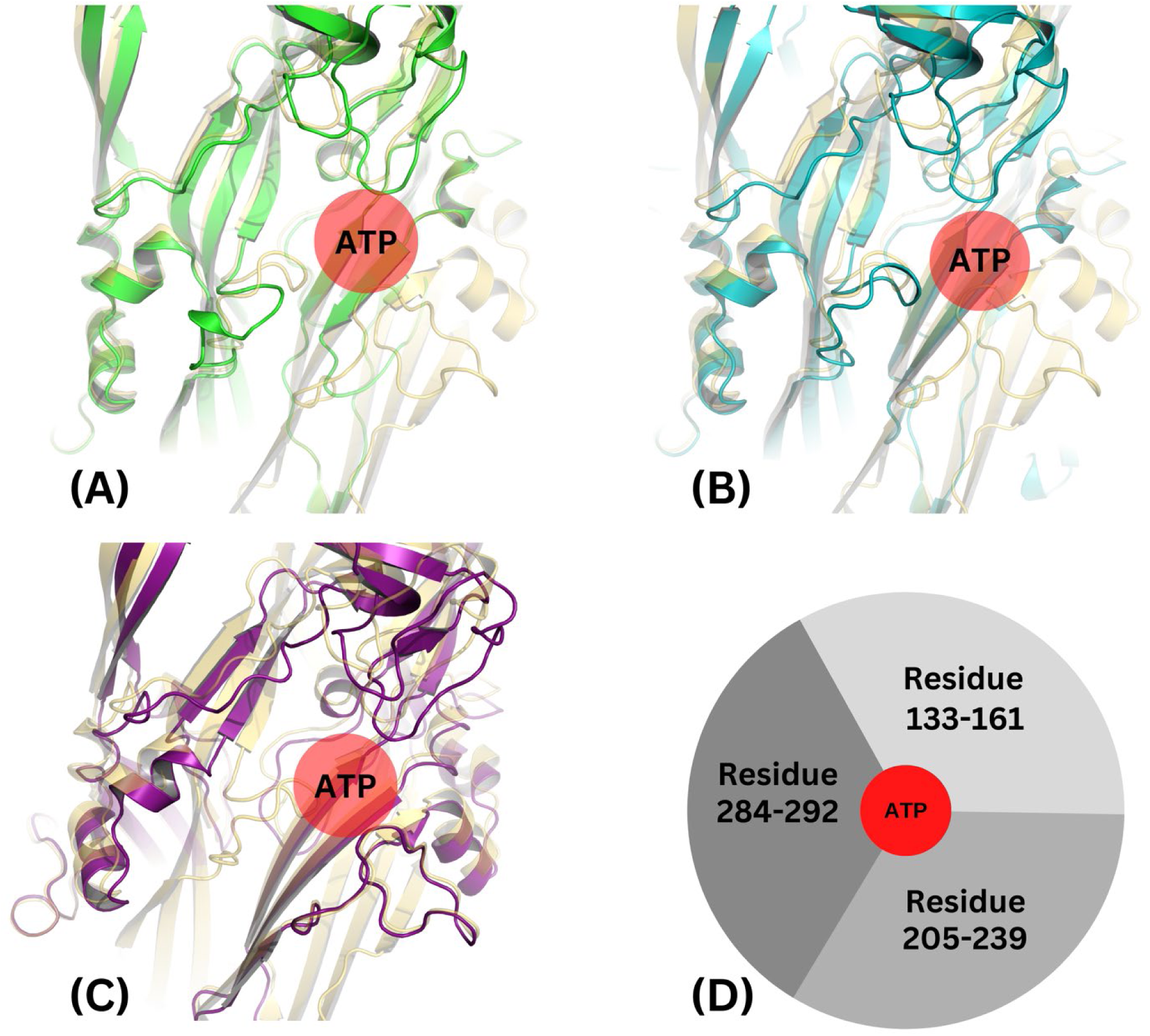
Model analysis of the sites of interest in the AF2M-generated P2X7R models. **(A)** ATP binding regions in the wild-type P2X7A (yellow) and P2X7E (green) subunits. **(B)** ATP binding regions in the wild-type P2X7A (yellow) and P2X7L (red) subunits. **(C)** ATP binding regions in the wild-type P2X7A (yellow) and P2X7J (dark purple) subunits. **(D)** Cartoon schematic of the ATP binding region demonstrating the three “lobes” of the region with their corresponding residue numbers, ATP (red) is shown in the centre.

**Table 5.** Residues within 6 Å of ATP in the main binding pocket. Residues shared by all four isoforms are shown in bold, residues shared by two or three isoforms are shown in italics.

| Wild-type P2X7A | P2X7E | P2X7L | P2X7J |
| --- | --- | --- | --- |
| **Lys66** | **Lys66** | **Lys66** | **Lys66** |
|  |  | Arg126 |  |
|  |  | Pro142 |  |
| **Gln143** | **Gln143** | **Gln143** | **Gln143** |
| **Thr189** | **Thr189** | **Thr189** | **Thr189** |
| *Leu191* | *Leu191* |  | *Leu191* |
| *Ile214* |  |  | *Ile214* |
| *Thr215* |  |  | *Thr215* |
|  |  |  | Cys216 |
| *Ile228* |  |  | *Ile228* |
| *Ser286* |  | *Ser286* |  |
| *Leu287* |  |  |  |
| *Tyr288* | *Tyr288* | *Tyr288* |  |

## Discussion

This study provides insight into the previously unknown structures of the human wild-type P2X7A, P2X7A/(A/E)/E, P2X7A/(A/B)/B, P2X7A/(A/J)/J, and P2X7A/(A/L)/L receptors derived using a deep learning predictive modelling method, AF2M. The models were validated and analysed using AF2M’s confidence measures and MD simulations, with the aim of providing templates for further P2X7R structure-function related research and drug discovery.

### Method validation

The computational modelling of protein structures has piqued the interest of researchers over the past three decades with biennial Critical Assessment of Structure Prediction (CASP) competitions run since 1994 (Moult, et al., 2014). The release of AlphaFold by the DeepMind group in 2021 enabled the prediction of protein structures to an experimental quality based on their amino acid sequence alone (Jumper, et al., 2021). The DeepMind group subsequently released an artificial intelligence modelling method, AF2M, that could predict the structures of protein complexes containing more than one subunit (Evans, et al., 2022). As AF2M is a novel protocol, there are limited published applications and validations of the method under different conditions and settings. Thus, in this study, we first investigated whether P2X7Rs, that are large receptor complexes, could be successfully modelled using AF2M. CASP14 competition criteria were used to validate the method (RMSD_experimental_, alignment score, and TM score) (Kryshtafovych, et al., 2014). The measures suggested the top-ranked AF2M-generated rat P2X7R model is a valid representation of the experimentally determined structure based on an RMSD_experimental_ value of less than 0.2 nm (Bordogna, et al., 2011), alignment score of less than 0.7 (Schrödinger Release_2022-3: Maestro, Schrödinger, LLC, New York, NY, 2021) and TM score of greater than 0.6 (Zhang and Skolnick, 2004).

The TM score (actual value) is 0.18 higher than the pTM score (predicted value). This finding supported Jumper, et al. (2021) who suggested the pTM is a conservative estimate of the TM score as it is based on the lower bounds of the pairwise matrix (a matrix used to identify the positional errors of the α-carbons in the structures) and so should be used as a comparator between the AF2M models through ranking the models from highest to lowest quality, rather than a definitive measure (Evans, et al., 2022, Jumper, et al., 2021). The RMSD_experimental_ was used to evaluate the ranking method of AF2M which uses the pTM and ipTM scores. The top-ranking model has the highest similarity to the experimental control, supporting the decision to select the top-ranked model for further study when considering the AF2M-generated heterotrimeric receptors.

As part of the method validation process, the addition of a custom template database was investigated. The AF2M models produced when a custom template database was used to inform the structure generation process were comparable to those produced when no template database was used. Originally, it was hypothesised that the custom templates may improve the predictions as they are integral to the original AlphaFold release (Jumper, et al., 2021). However, based on the results the quality of the AF2M predictions was likely dependent on the quality of the MSA representations rather than the templates as supported by Evans, et al. (2022).

### Heterotrimeric receptors

The structures of P2X7Rs that incorporate splice variants in combination with one or two wild-type P2X7A subunits were modelled. These structures present a challenge to traditional homology modelling methods as they are missing large domains inside the protein sequence. Deep learning methods such as AF2M can overcome this limitation of homology modelling. These computational models can be used to aid the elucidation of further cryo-EM or X-ray crystallography structures as highlighted by Kryshtafovych, et al. (2021) who used computational models as a template for molecular replacement in X-ray crystallography, and backbone tracing in cryo-EM (Kryshtafovych, et al., 2021). Visual inspection of the structures, AF2M’s confidence measures, and MD simulation outputs (RMSD_MD_ and RMSF) were used to determine the quality of the AF2M models. These measures were used as no experimentally obtained structures are available for the splice variants.

Overall, the top-ranking models were predicted with high confidence in relation to the pLDDT output, suggesting the models are good predictions of the protein backbone (Jumper, et al., 2021). The largest drop in confidence from residues 433 to 472 (a loop region in the CTD) in the wild-type P2X7A subunits and P2X7L splice variants can be attributed to firstly, the structural irregularity of loop regions in comparison to more stable α helices and β strands (Fiser, et al., 2000), and secondly, a reduction in the number and sequence similarity of the MSAs (which are required to inform the structure generation process (Mirdita, et al., 2022)) for the CTD due to its unique structure which is not observed in any of the other six P2X receptors (Kopp, et al., 2019). The reduction in per-residue confidence in the residues near the spliced-out sites in the splice variants can be explained by the reduction in inter-residue contacts in these regions that are key to informing the structure generation process. This is supported by Jumper, et al. (2021) who suggested that higher numbers of inter-residue contacts are linked with higher prediction confidence scores.

MD simulations were used as an alternate method to validate the AF2M structures in a membrane system, a validation method that has previously been used in the literature to confirm the stability of models (Khare, et al., 2022, Radom, et al., 2018, Soni, et al., 2014). The consistency between the predicted models and the experimental control, whereby all the receptors stabilised and reported similar flexible regions, suggest all the models are stable structures. Flexible residues are located in the loop regions of the receptor that are exposed to the extracellular and intracellular environment, flexibility which is expected due to the reduced structural integrity of loop regions (Fiser, et al., 2000).

The intracellular ATD and CTD in the models are of particular interest. P2X7Rs are unique compared to other P2X receptors with the presence of a large CTD that contains key functional regions including the Cys-rich domain, tumour necrosis factor receptor 1 death domain, lipopolysaccharide domain, actin filament binding domain, sarcoma tyrosine kinase homology 3 binding domain, and trafficking domain (De Salis, et al., 2022). It is therefore of interest that the P2X7B, P2X7E, and P2X7J splice variants are missing these CTD regions. The RMSF results revealed that the ATD and CTD in the wild-type P2X7A subunits are significantly more flexible in the P2X7A/B/B, P2X7A/E/E, and P2X7A/J/J receptors, likely as the intracellular CTDs are missing in the P2X7B, P2X7E, and P2X7J splice variants so there are less structures limiting the movement of the termini. In turn, the higher flexibility of the intracellular termini of these receptors that are composed of two splice variants translates to a higher overall flexibility over time as observed in the RMSD_MD_ values. Further studies are required to determine the impact of these findings in health and disease.

Study of the ATP binding sites is also crucial to understanding the function of the heterotrimeric receptors. Residues within the original binding pockets in the wild-type P2X7A receptors were found to be absent at the interface between the P2X7E/P2X7J/P2X7L and P2X7A subunits, and new residue contacts extended into the altered space from nearby loops. Alterations in the splice variants did not appear to affect the binding site conformations in the P2X7A subunits of the heterotrimeric receptors. These results suggest P2X7A/A/E, P2X7A/A/J, and P2X7A/A/L receptors could have two functional ATP binding sites while P2X7A/E/E, P2X7A/J/J, and P2X7A/L/L receptors could have one functional ATP binding site. The modelling of the ATP binding sites in the P2X7E and P2X7L heterotrimers aligns with the latest literature whereby Skarratt, et al. (2020) hypothesised the P2X7E and P2X7L subunits lack the ATP binding sites that are found in wild-type P2X7A subunits. Further work is required to support this observation.

### Limitations

It is important to acknowledge that the structures are predictions rather than experimental structures and therefore should be continuously validated as new experimental data becomes available (Evans, et al., 2022). Further refinement of the models may be possible using some of the recent developments presented at the CASP15 competition such as D-I-TASSER (Zheng, et al., 2021) and trROSETTA (Du, et al., 2021, Wang, et al., 2022, Su, et al., 2021, Yang, et al., 2020). These methods consider different approaches and implementations of the standard AF2M.

For the MD simulations we used a simple membrane (composed of POPC and cholesterol molecules) rather than a more complex membrane system that reflects the composition of human epithelial membranes. The simpler membrane was deemed appropriate for this study as the receptor rather than the membrane characteristics were being studied. Further studies should investigate the receptor stability and interactions with the membrane environment in different membrane systems to account for the variability in membrane composition throughout the human body (Wilson, et al., 2021) as well as the crucial modulatory effect of membrane cholesterol on P2X7R function (Robinson, et al., 2014).

### Conclusion and future direction

In the present study we have evaluated the suitability of the deep learning approach AF2M in predicting the structure of large purinergic receptor protein complexes, more specifically the P2X7R. The developed protocol was used to generate models of P2X7R complexes incorporating alternatively spliced subunits together with wild type subunits in 1:2 and 2:1 stoichiometric ratios. The models were validated using AF2M confidence metrics and further supported by MD simulation results which showed that the generated models were overall stable over time.

The present study provides the first insight into the structure of P2X7Rs that incorporate alternatively spliced subunits. Precision medicine is an emerging model for treating patients with conditions that are clinically and biologically heterogeneous and therefore require individualized treatment. We have developed a model pipeline for analysis of binding of known antagonists and novel molecules to P2X7R alternatively spliced isoforms that can be translated into treatments for cancers, inflammatory conditions, trauma, and neurodegenerative diseases.

## Supporting information

Supplementary_Models

## Acknowledgements

This research was supported by the Australian Government’s National Collaborative Research Infrastructure Strategy, with access to computational resources provided by the National Computing Infrastructure through the National Computational Merit Allocation Scheme (NCMAS-2022-154). Furthermore, the authors would like to acknowledge the technical assistance provided by the Sydney Informatics Hub, a Core Research Facility of the University of Sydney.

Jake Zheng Chen is funded by the Australian Government Research Training Program Scholarship.

## Author contributions

Sophie K. F. De Salis participated in the research design, conducted experiments, performed data analysis, and wrote the manuscript. Jake Zheng Chen participated in the research design, conducted experiments, performed data analysis, and wrote the manuscript. Kristen K. Skarratt participated in the preparation of the manuscript and research design. Stephen J. Fuller and Thomas Balle participated in the research design, data analysis and supervised the writing of the manuscript. All authors were involved in the revision of the manuscript. All authors have approved this manuscript.

## References

Abraham, M. J., Murtola, T., Schulz, R., Páll, S., Smith, J. C., Hess, B. and Lindahl, E. 2015. GROMACS: High performance molecular simulations through multi-level parallelism from laptops to supercomputers. SoftwareX, 1-2:19–25. doi:10.1016/j.softx.2015.06.001.

Adinolfi, E., Callegari, M. G., Ferrari, D., Bolognesi, C., Minelli, M., Wieckowski, M. R., Pinton, P., Rizzuto, R. and Di Virgilio, F. 2005. Basal activation of the P2X7 ATP receptor elevates mitochondrial calcium and potential, increases cellular ATP levels, and promotes serum-independent growth. Mol Biol Cell, 16:3260–3272. doi:10.1091/mbc.e04-11-1025.

Adinolfi, E., Cirillo, M., Woltersdorf, R., Falzoni, S., Chiozzi, P., Pellegatti, P., Callegari, M. G., Sandonà, D., Markwardt, F., Schmalzing, G. and Di Virgilio, F. 2010. Trophic activity of a naturally occurring truncated isoform of the P2X7 receptor. FASEB J, 24:3393–3404. doi:10.1096/fj.09-153601.

Bahari, G., Tabasi, F., Hashemi, M., Zakeri, Z. and Taheri, M. 2021. Association of P2X7 receptor genetic polymorphisms and expression with rheumatoid arthritis susceptibility in a sample of the Iranian population: a case-control study. Clin Rheumatol, 40:3115–3126. doi:10.1007/s10067-021-05645-3.

Berendsen, H. J. C., Postma, J. P. M., van Gunsteren, W. F., DiNola, A. and Haak, J. R. 1984. Molecular dynamics with coupling to an external bath. J Chem Phys 81:3684–3690. doi:10.1063/1.448118.

Berendsen, H. J. C., van der Spoel, D. and van Drunen, R. 1995. GROMACS: A message-passing parallel molecular dynamics implementation. Comput Phys Commun, 91:43–56. doi:10.1016/0010-4655(95)00042-E.

Bordogna, A., Pandini, A. and Bonati, L. 2011. Predicting the accuracy of protein-ligand docking on homology models. J Comput Chem, 32:81–98. doi:10.1002/jcc.21601.

Brooks, B. R., Brooks, C. L., 3rd, Mackerell, A. D., Jr., Nilsson, L., Petrella, R. J., Roux, B., Won, Y., Archontis, G., Bartels, C., Boresch, S., Caflisch, A., Caves, L., Cui, Q., Dinner, A. R., Feig, M., Fischer, S., Gao, J., Hodoscek, M., Im, W., Kuczera, K., Lazaridis, T., Ma, J., Ovchinnikov, V., Paci, E., Pastor, R. W., Post, C. B., Pu, J. Z., Schaefer, M., Tidor, B., Venable, R. M., Woodcock, H. L., Wu, X., Yang, W., York, D. M. and Karplus, M. 2009. CHARMM: the biomolecular simulation program. J Comput Chem, 30:1545–1614. doi:10.1002/jcc.21287.

Case, D. A., Cheatham, T. E., 3rd, Darden, T., Gohlke, H., Luo, R., Merz, K. M., Jr., Onufriev, A., Simmerling, C., Wang, B. and Woods, R. J. 2005. The Amber biomolecular simulation programs. J Comput Chem, 26:1668–1688. doi:10.1002/jcc.20290.

Cauchy, A. 1847. Méthode générale pour la résolution des systèmes d’équations simultanées. C R Acad Sci, 25:536–538.

Cheewatrakoolpong, B., Gilchrest, H., Anthes, J. C. and Greenfeder, S. 2005. Identification and characterization of splice variants of the human P2X_7_ ATP channel. Biochem Biophys Res Commun, 332:17–27. doi:10.1016/j.bbrc.2005.04.087.

Clark, K., Karsch-Mizrachi, I., Lipman, D. J., Ostell, J. and Sayers, E. W. 2016. GenBank. Nucleic Acids Res, 44:D67–D72. doi:10.1093/nar/gkv1276.

Du, Z., Su, H., Wang, W., Ye, L., Wei, H., Peng, Z., Anishchenko, I., Baker, D. and Yang, J. 2021. The trRosetta server for fast and accurate protein structure prediction. Nature Protocols, 16:5634–5651. doi:10.1038/s41596-021-00628-9.

Essmann, U., Perera, L., Berkowitz, M. L., Darden, T., Lee, H. and Pedersen, L. G. 1995. A smooth particle mesh Ewald method. J Chem Phys, 103:8577–8593. doi:10.1063/1.470117.

Evans, R., O’Neill, M., Pritzel, A., Antropova, N., Senior, A., Green, T., Žídek, A., Bates, R., Blackwell, S., Yim, J., Ronneberger, O., Bodenstein, S., Zielinski, M., Bridgland, A., Potapenko, A., Cowie, A., Tunyasuvunakool, K., Jain, R., Clancy, E., Kohli, P., Jumper, J. and Hassabis, D. 2022. Protein complex prediction with AlphaFold-Multimer. bioRxiv. doi:10.1101/2021.10.04.463034.

Feng, Y. H., Li, X., Wang, L., Zhou, L. and Gorodeski, G. I. 2006. A truncated P2X7 receptor variant (P2X_7-j_) endogenously expressed in cervical cancer cells antagonizes the full-length P2X7 receptor through hetero-oligomerization. J Biol Chem, 281:17228–17237. doi:10.1074/jbc.M602999200.

Fiser, A., Do, R. K. and Sali, A. 2000. Modeling of loops in protein structures. Protein Sci, 9:1753–1773. doi:10.1110/ps.9.9.1753.

Fryatt, A. G., Dayl, S., Stavrou, A., Schmid, R. and Evans, R. J. 2019. Organization of ATP-gated P2X1 receptor intracellular termini in apo and desensitized states. J Gen Physiol, 151:146–155. doi:10.1085/jgp.201812108.

Giuliani, A. L., Colognesi, D., Ricco, T., Roncato, C., Capece, M., Amoroso, F., Wang, Q. G., De Marchi, E., Gartland, A., Di Virgilio, F. and Adinolfi, E. 2014. Trophic activity of human P2X7 receptor isoforms A and B in osteosarcoma. PLoS One, 9:e107224. doi:10.1371/journal.pone.0107224.

Hess, B., Bekker, H., Berendsen, H. J. C. and Fraaije, J. G. E. M. 1997. LINCS: A linear constraint solver for molecular simulations. J Comput Chem, 18:1463–1472. doi:10.1002/(SICI)1096-987X(199709)18:12<1463::AID-JCC4>3.0.CO;2-H.

Hess, B., Kutzner, C., van der Spoel, D. and Lindahl, E. 2008. GROMACS 4: algorithms for highly efficient, load-balanced, and scalable molecular simulation. J Chem Theory Comput, 4:435–447. doi:10.1021/ct700301q.

Hilbert, M., Böhm, G. and Jaenicke, R. 1993. Structural relationships of homologous proteins as a fundamental principle in homology modeling. Proteins, 17:138–151. doi:10.1002/prot.340170204.

Hoover, W. G. 1985. Canonical dynamics: Equilibrium phase-space distributions. Phys Rev A, 31:1695–1697. doi:10.1103/PhysRevA.31.1695.

Illes, P., Müller, C. E., Jacobson, K. A., Grutter, T., Nicke, A., Fountain, S. J., Kennedy, C., Schmalzing, G., Jarvis, M. F., Stojilkovic, S. S., King, B. F. and Di Virgilio, F. 2021. Update of P2X receptor properties and their pharmacology: IUPHAR Review 30. Br J Pharmacol, 178:489–514. doi:10.1111/bph.15299.

Ingólfsson, H. I., Melo, M. N., van Eerden, F. J., Arnarez, C., Lopez, C. A., Wassenaar, T. A., Periole, X., de Vries, A. H., Tieleman, D. P. and Marrink, S. J. 2014. Lipid organization of the plasma membrane. J Am Chem Soc, 136:14554–14559. doi:10.1021/ja507832e.

Jo, S., Kim, T. and Im, W. 2007. Automated builder and database of protein/membrane complexes for molecular dynamics simulations. PLoS One, 2:e880. doi:10.1371/journal.pone.0000880.

Jo, S., Kim, T., Iyer, V. G. and Im, W. 2008. CHARMM-GUI: a web-based graphical user interface for CHARMM. J Comput Chem, 29:1859–1865. doi:10.1002/jcc.20945.

Jo, S., Lim, J. B., Klauda, J. B. and Im, W. 2009. CHARMM-GUI membrane builder for mixed bilayers and its application to yeast membranes. Biophys J, 97:50–58. doi:10.1016/j.bpj.2009.04.013.

Jumper, J., Evans, R., Pritzel, A., Green, T., Figurnov, M., Ronneberger, O., Tunyasuvunakool, K., Bates, R., Žídek, A., Potapenko, A., Bridgland, A., Meyer, C., Kohl, S. A. A., Ballard, A. J., Cowie, A., Romera-Paredes, B., Nikolov, S., Jain, R., Adler, J., Back, T., Petersen, S., Reiman, D., Clancy, E., Zielinski, M., Steinegger, M., Pacholska, M., Berghammer, T., Bodenstein, S., Silver, D., Vinyals, O., Senior, A. W., Kavukcuoglu, K., Kohli, P. and Hassabis, D. 2021. Highly accurate protein structure prediction with AlphaFold. Nature, 596:583–589. doi:10.1038/s41586-021-03819-2.

Karasawa, A. and Kawate, T. 2016. Structural basis for subtype-specific inhibition of the P2X7 receptor. eLife, 5:e22153. doi:10.7554/eLife.22153.

Kawate, T., Michel, J. C., Birdsong, W. T. and Gouaux, E. 2009. Crystal structure of the ATP-gated P2X_4_ ion channel in the closed state. Nature, 460:592–598. doi:10.1038/nature08198.

Keystone, E. C., Wang, M. M., Layton, M., Hollis, S. and McInnes, I. B. 2012. Clinical evaluation of the efficacy of the P2X7 purinergic receptor antagonist AZD9056 on the signs and symptoms of rheumatoid arthritis in patients with active disease despite treatment with methotrexate or sulphasalazine. Ann Rheum Dis, 71:1630–1635. doi:10.1136/annrheumdis-2011-143578.

Khare, N., Maheshwari, S. K., Rizvi, S. M. D., Albadrani, H. M., Alsagaby, S. A., Alturaiki, W., Iqbal, D., Zia, Q., Villa, C., Jha, S. K., Jha, N. K. and Jha, A. K. 2022. Homology modelling, molecular docking and molecular dynamics simulation studies of CALMH1 against secondary metabolites of bauhinia variegata to treat Alzheimer’s disease. Brain Sci, 12:770. doi:10.3390/brainsci12060770.

Kopp, R., Krautloher, A., Ramírez-Fernández, A. and Nicke, A. 2019. P2X7 interactions and signaling - making head or tail of it. Front Mol Neurosci, 12:183. doi:10.3389/fnmol.2019.00183.

Kryshtafovych, A., Monastyrskyy, B. and Fidelis, K. 2014. CASP prediction center infrastructure and evaluation measures in CASP10 and CASP ROLL. Proteins, 82:7–13. doi:10.1002/prot.24399.

Kryshtafovych, A., Moult, J., Albrecht, R., Chang, G. A., Chao, K., Fraser, A., Greenfield, J., Hartmann, M. D., Herzberg, O., Josts, I., Leiman, P. G., Linden, S. B., Lupas, A. N., Nelson, D. C., Rees, S. D., Shang, X., Sokolova, M. L. and Tidow, H. 2021. Computational models in the service of X-ray and cryo-electron microscopy structure determination. Proteins, 89:1633–1646. doi:10.1002/prot.26223.

Lee, J., Cheng, X., Swails, J. M., Yeom, M. S., Eastman, P. K., Lemkul, J. A., Wei, S., Buckner, J., Jeong, J. C., Qi, Y., Jo, S., Pande, V. S., Case, D. A., Brooks, C. L., MacKerell, A. D., Klauda, J. B. and Im, W. 2016. CHARMM-GUI Input Generator for NAMD, GROMACS, AMBER, OpenMM, and CHARMM/OpenMM Simulations Using the CHARMM36 Additive Force Field. J Chem Theory Comput, 12:405–413. doi:10.1021/acs.jctc.5b00935.

Lee, J., Patel, D. S., Ståhle, J., Park, S. J., Kern, N. R., Kim, S., Lee, J., Cheng, X., Valvano, M. A., Holst, O., Knirel, Y. A., Qi, Y., Jo, S., Klauda, J. B., Widmalm, G. and Im, W. 2019. CHARMM-GUI membrane builder for complex biological membrane simulations with glycolipids and lipoglycans. J Chem Theory Comput, 15:775–786. doi:10.1021/acs.jctc.8b01066.

Li, M., Toombes, G. E., Silberberg, S. D. and Swartz, K. J. 2015. Physical basis of apparent pore dilation of ATP-activated P2X receptor channels. Nat neurosci, 18:1577–1583. doi:10.1038/nn.4120.

Lindahl, E., Abraham, M. J., Hess, B. and van der Spoel, D. 2021. GROMACS 2021.4 source code. Zenodo. doi:10.5281/zenodo.5636567.

Lindahl, E., Hess, B. and van der Spoel, D. 2001. GROMACS 3.0: a package for molecular simulation and trajectory analysis. J Mol Model, 7:306–317. doi:10.1007/s008940100045.

Liu, X., Zhao, Z., Ji, R., Zhu, J., Sui, Q. Q., Knight, G. E., Burnstock, G., He, C., Yuan, H. and Xiang, Z. 2017. Inhibition of P2X7 receptors improves outcomes after traumatic brain injury in rats. Purinergic Signal, 13:529–544. doi:10.1007/s11302-017-9579-y.

Lomize, M. A., Pogozheva, I. D., Joo, H., Mosberg, H. I. and Lomize, A. L. 2012. OPM database and PPM web server: resources for positioning of proteins in membranes. Nucleic Acids Res, 40:D370–D376. doi:10.1093/nar/gkr703.

Lu, C., Wu, C., Ghoreishi, D., Chen, W., Wang, L., Damm, W., Ross, G. A., Dahlgren, M. K., Russell, E., Von Bargen, C. D., Abel, R., Friesner, R. A. and Harder, E. D. 2021. OPLS4: improving force field accuracy on challenging regimes of chemical space. J Chem Theory Comput, 17:4291–4300. doi:10.1021/acs.jctc.1c00302.

Martin, E., Amar, M., Dalle, C., Youssef, I., Boucher, C., Le Duigou, C., Brückner, M., Prigent, A., Sazdovitch, V., Halle, A., Kanellopoulos, J. M., Fontaine, B., Delatour, B. and Delarasse, C. 2019. New role of P2X7 receptor in an Alzheimer’s disease mouse model. Mol Psychiatry, 24:108–125. doi:10.1038/s41380-018-0108-3.

McCarthy, A. E., Yoshioka, C. and Mansoor, S. E. 2019. Full-length P2X_7_ structures reveal how palmitoylation prevents channel desensitization. Cell, 179:659–670.e613. doi:10.1016/j.cell.2019.09.017.

Mirdita, M., Schütze, K., Moriwaki, Y., Heo, L., Ovchinnikov, S. and Steinegger, M. 2022. ColabFold: making protein folding accessible to all. Nat Methods, 19:679–682. doi:10.1038/s41592-022-01488-1.

Mirdita, M., Steinegger, M., Breitwieser, F., Söding, J. and Levy Karin, E. 2021. Fast and sensitive taxonomic assignment to metagenomic contigs. Bioinformatics, 37:3029–3031. doi:10.1093/bioinformatics/btab184.

Mirdita, M., Steinegger, M. and Söding, J. 2019. MMseqs2 desktop and local web server app for fast, interactive sequence searches. Bioinformatics, 35:2856–2858. doi:10.1093/bioinformatics/bty1057.

Moult, J., Fidelis, K., Kryshtafovych, A., Schwede, T. and Tramontano, A. 2014. Critical assessment of methods of protein structure prediction (CASP)-round x. Proteins, 82:1–6. doi:10.1002/prot.24452.

Nosé, S. 1984. A molecular dynamics method for simulations in the canonical ensemble. Mol Phys, 52:255–268. doi:10.1080/00268978400101201.

Notomi, S., Hisatomi, T., Kanemaru, T., Takeda, A., Ikeda, Y., Enaida, H., Kroemer, G. and Ishibashi, T. 2011. Critical involvement of extracellular ATP acting on P2RX7 purinergic receptors in photoreceptor cell death. Am J Pathol, 179:2798–2809. doi:10.1016/j.ajpath.2011.08.035.

Olsson, M. H. M., Søndergaard, C. R., Rostkowski, M. and Jensen, J. H. 2011. PROPKA3: consistent treatment of internal and surface residues in empirical pKa predictions. J Chem Theory Comput, 7:525–537. doi:10.1021/ct100578z.

Páll, S. and Hess, B. 2013. A flexible algorithm for calculating pair interactions on SIMD architectures. Comput Phys Commun, 184:2641–2650. doi:10.1016/j.cpc.2013.06.003.

Papadopoulos, J. S. and Agarwala, R. 2007. COBALT: constraint-based alignment tool for multiple protein sequences. Bioinformatics, 23:1073–1079. doi:10.1093/bioinformatics/btm076.

Parrinello, M. and Rahman, A. 1981. Polymorphic transitions in single crystals: A new molecular dynamics method. J Appl Phys, 52:7182–7190. doi:10.1063/1.328693.

Pegoraro, A., De Marchi, E. and Adinolfi, E. 2021. P2X7 variants in oncogenesis. Cells, 10:189. doi:10.3390/cells10010189.

Pegoraro, A., Orioli, E., De Marchi, E., Salvestrini, V., Milani, A., Di Virgilio, F., Curti, A. and Adinolfi, E. 2020. Differential sensitivity of acute myeloid leukemia cells to daunorubicin depends on P2X7A versus P2X7B receptor expression. Cell Death Dis, 11:876. doi:10.1038/s41419-020-03058-9.

Pronk, S., Páll, S., Schulz, R., Larsson, P., Bjelkmar, P., Apostolov, R., Shirts, M. R., Smith, J. C., Kasson, P. M., van der Spoel, D., Hess, B. and Lindahl, E. 2013. GROMACS 4.5: a high-throughput and highly parallel open source molecular simulation toolkit. Bioinformatics, 29:845–854. doi:10.1093/bioinformatics/btt055.

Radom, F., Plückthun, A. and Paci, E. 2018. Assessment of *ab initio* models of protein complexes by molecular dynamics. PLoS Comput Biol, 14:e1006182. doi:10.1371/journal.pcbi.1006182.

Rassendren, F., Buell, G. N., Virginio, C., Collo, G., North, R. A. and Surprenant, A. 1997. The permeabilizing ATP receptor, P2X7. Cloning and expression of a human cDNA. J Biol Chem, 272:5482–5486. doi:10.1074/jbc.272.9.5482.

Riedel, T., Schmalzing, G. and Markwardt, F. 2007. Influence of extracellular monovalent cations on pore and gating properties of P2X7 receptor-operated single-channel currents. Biophys J, 93:846–858. doi:10.1529/biophysj.106.103614.

Robinson, L. E., Shridar, M., Smith, P. and Murrell-Lagnado, R. D. 2014. Plasma membrane cholesterol as a regulator of human and rodent P2X7 receptor activation and sensitization. J Biol Chem, 289:31983–31994. doi:10.1074/jbc.M114.574699.

Salomon-Ferrer, R., Case, D. A. and Walker, R. C. 2013. An overview of the Amber biomolecular simulation package. WIREs Comput Mol Sci, 3:198–210. doi:10.1002/wcms.1121.

Sastry, G. M., Adzhigirey, M., Day, T., Annabhimoju, R. and Sherman, W. 2013. Protein and ligand preparation: parameters, protocols, and influence on virtual screening enrichments. J Comput Aided Mol Des, 27:221–234. doi:10.1007/s10822-013-9644-8.

Skarratt, K. K., Gu, B. J., Lovelace, M. D., Milligan, C. J., Stokes, L., Glover, R., Petrou, S., Wiley, J. S. and Fuller, S. J. 2020. A P2RX7 single nucleotide polymorphism haplotype promotes exon 7 and 8 skipping and disrupts receptor function. FASEB J 34:3884–3901. doi:10.1096/fj.201901198RR.

Søndergaard, C. R., Olsson, M. H. M., Rostkowski, M. and Jensen, J. H. 2011. Improved treatment of ligands and coupling effects in empirical calculation and rationalization of pKa values. J Chem Theory Comput, 7:2284–2295. doi:10.1021/ct200133y.

Soni, S., Tyagi, C., Grover, A. and Goswami, S. K. 2014. Molecular modeling and molecular dynamics simulations based structural analysis of the SG2NA protein variants. BMC Res Notes, 7:446. doi:10.1186/1756-0500-7-446.

Steinegger, M. and Söding, J. 2018. Clustering huge protein sequence sets in linear time. Nat Commun, 9:2542. doi:10.1038/s41467-018-04964-5.

Steinegger, M. and Söding, J. 2017. MMseqs2 enables sensitive protein sequence searching for the analysis of massive data sets. Nat Biotechnol, 35:1026–1028. doi:10.1038/nbt.3988.

Stock, T. C., Bloom, B. J., Wei, N., Ishaq, S., Park, W., Wang, X., Gupta, P. and Mebus, C. A. 2012. Efficacy and safety of CE-224,535, an antagonist of P2X7 receptor, in treatment of patients with rheumatoid arthritis inadequately controlled by methotrexate. J Rheumatol, 39:720–727. doi:10.3899/jrheum.110874.

Su, H., Wang, W., Du, Z., Peng, Z., Gao, S. H., Cheng, M. M. and Yang, J. 2021. Improved protein structure prediction using a new multi-scale network and homologous templates. Adv Sci (Weinh), 8:e2102592. doi:10.1002/advs.202102592.

The UniProt, C. 2021. UniProt: the universal protein knowledgebase in 2021. Nucleic Acids Res, 49:D480–D489. doi:10.1093/nar/gkaa1100.

Ulrich, H., Ratajczak, M. Z., Schneider, G., Adinolfi, E., Orioli, E., Ferrazoli, E. G., Glaser, T., Corrêa-Velloso, J., Martins, P. C. M., Coutinho, F., Santos, A. P. J., Pillat, M. M., Sack, U. and Lameu, C. 2018. Kinin and purine signaling contributes to neuroblastoma metastasis. Front Pharmacol, 9. doi:10.3389/fphar.2018.00500.

Van Der Spoel, D., Lindahl, E., Hess, B., Groenhof, G., Mark, A. E. and Berendsen, H. J. 2005. GROMACS: fast, flexible, and free. J Comput Chem, 26:1701–1718. doi:10.1002/jcc.20291.

Virginio, C., MacKenzie, A., North, R. A. and Surprenant, A. 1999. Kinetics of cell lysis, dye uptake and permeability changes in cells expressing the rat P2X7 receptor. J Physiol, 519:335–346. doi:10.1111/j.1469-7793.1999.0335m.x.

Wang, W., Peng, Z. and Yang, J. 2022. Single-sequence protein structure prediction using supervised transformer protein language models. bioRxiv:2022.2001.2015.476476. doi:10.1101/2022.01.15.476476.

Wilson, K. A., Fairweather, S. J., MacDermott-Opeskin, H. I., Wang, L., Morris, R. A. and O’Mara, M. L. 2021. The role of plasmalogens, Forssman lipids, and sphingolipid hydroxylation in modulating the biophysical properties of the epithelial plasma membrane. J Chem Phys, 154:095101. doi:10.1063/5.0040887.

Wu, E. L., Cheng, X., Jo, S., Rui, H., Song, K. C., Dávila-Contreras, E. M., Qi, Y., Lee, J., Monje-Galvan, V., Venable, R. M., Klauda, J. B. and Im, W. 2014. CHARMM-GUI membrane builder toward realistic biological membrane simulations. J Comput Chem, 35:1997–2004. doi:10.1002/jcc.23702.

Xu, J. and Zhang, Y. 2010. How significant is a protein structure similarity with TM-score = 0.5? Bioinformatics, 26:889–895. doi:10.1093/bioinformatics/btq066.

Yang, J., Anishchenko, I., Park, H., Peng, Z., Ovchinnikov, S. and Baker, D. 2020. Improved protein structure prediction using predicted interresidue orientations. Proc Natl Acad Sci U S A, 117:1496–1503. doi:10.1073/pnas.1914677117.

Zhang, X., Zheng, G., Ma, X., Yang, Y., Li, G., Rao, Q., Nie, K. and Wu, K. 2004. Expression of P2X7 in human hematopoietic cell lines and leukemia patients. Leuk Res, 28:1313–1322. doi:10.1016/j.leukres.2004.04.001.

Zhang, Y. and Skolnick, J. 2004. Scoring function for automated assessment of protein structure template quality. Proteins, 57:702–710. doi:10.1002/prot.20264.

Zheng, W., Zhang, C., Li, Y., Pearce, R., Bell, E. W. and Zhang, Y. 2021. Folding non-homologous proteins by coupling deep-learning contact maps with I-TASSER assembly simulations. Cell Rep Methods, 1. doi:10.1016/j.crmeth.2021.100014.

